# Diptera-specific Daedalus controls Zucchini endonucleolysis in piRNA biogenesis independent of exonucleases

**DOI:** 10.1101/2024.06.02.596995

**Authors:** Yuica Koga, Shigeki Hirakata, Mayu Negishi, Hiroya Yamazaki, Mikiko C. Siomi

## Abstract

piRNAs protect germline genomes and maintain fertility by repressing transposons. Daedalus and Gasz act together as a mitochondrial scaffold for Armitage, a necessary factor for Zucchini-dependent piRNA processing. However, the mechanism underlying this function remains unclear. Here, we find that the roles of Daedalus and Gasz in this process are distinct, although both are necessary: Daedalus physically interacts with Armitage, whereas Gasz supports Daedalus to maintain its function. Daedalus binds to Armitage through two distinct regions, an extended coiled-coil identified in this study, and a SAM. The former tethers Armitage to mitochondria, while the latter controls Zucchini endonucleolysis to define the length of piRNAs in an exonuclease-independent manner. piRNAs produced in the absence of Daedalus SAM do not exhibit full transposon silencing functionality. Daedalus is Diptera specific. Unlike *Drosophila* and mosquitoes, evolutionarily more complex species, such as mice, rely on exonucleases after Zucchini processing to specify the length of piRNAs.

## Introduction

In *Drosophila* ovarian somatic cells (OSCs), PIWI-interacting RNAs (piRNAs) associate with one of the PIWI members, Piwi, in the cytoplasm to form the piRNA-induced silencing complex (piRISC).^1–4^ piRISC then co-transcriptionally represses transposons in the nucleus. The vast majority of piRNAs in OSCs originate from the piRNA cluster, *flamenco* (*flam*).^5^ The *flam* locus contains numerous transposon remnants, most of which have an orientation opposite to the direction of transcription. Therefore, piRNAs can naturally hybridize to target mRNAs through RNA–RNA base pairing.^5,6^ After nuclear processing, including splicing,^7^ the *flam* transcripts are exported to the cytoplasm where they bind to female sterile (1) Yb (Yb) via *cis-*elements embedded in the RNAs. This interaction induces the assembly of non-membranous organelle Yb bodies through phase separation, which accelerates subsequent steps in the piRNA processing.^8,9^

At this point, the *flam* transcripts are coarsely digested by an unknown factor, generating numerous 5′ ends. Piwi binds one-to-one to these 5′ ends via the MID domain and becomes the Piwi-piRISC precursor (pre-Piwi-piRISC).^10,11^ Concomitantly, the RNA helicase Armitage (Armi) binds to the complex, allowing pre-Piwi-piRISC to exit Yb bodies and move to the mitochondrial surface where the endonuclease Zucchini (Zuc) awaits to process the complex.^12^ Zuc has a transmembrane (TM) region, which localizes Zuc to the outer membrane of mitochondria.^13–16^ After Zuc RNA cleavage releases mature Piwi-piRISC from pre-Piwi-piRISC, nascent Piwi binds to the remaining piRNA precursor through its 5′ end, generating a new pre-Piwi-piRISC in Yb body-independent and Zuc-dependent manners.^17,18^ This mechanism of sequential generation of Piwi-piRISC from a single piRNA precursor is known as phasing.^17,18^ Piwi-piRISC is then transported to the nucleus,^19^ where it associates with nascent transposon transcripts and suppresses transcription in concert with multiple cofactors, such as Gtsf1/Asterix and Maelstrom.^20–27^

An RNAi-based gene screen performed in *Drosophila* ovaries identified many factors acting in the piRNA pathway, including Gasz and Daedalus (Daed).^28^ Both proteins have a TM, like Zuc, and they form a heteromeric complex that acts in concert with Armi to support Zuc cleavage in piRNA maturation.^12,29,30^ However, the molecular details of this process, especially the function of Daed, remain unclear.

In this study, we found that Daed interacts with Armi through two distinct regions, the extended coiled-coil (eCC) identified in this study, and the sterile α motif (SAM). We also show that Daed interacts with Gasz by forming a helical barrel using the two α-helixes present in each of Daed and Gasz. Through this interaction, Daed stabilizes Gasz, but not *vice versa*. Furthermore, we found that the Armi–Daed interaction through Daed eCC tethers Armi to mitochondria, while the Armi–Daed interaction through SAM controls Zuc endonucleolysis to specify the length of piRNAs in an exonuclease-independent manner. piRNAs produced in the absence of Daed SAM became atypically long and their function in transposon silencing was attenuated. Evolutionarily more complex species, such as mice, possess exonuclease, PNLDC1, but not Daed. Mouse piRNAs, which are atypically elongated in the absence of PNLDC1, do not exhibit full functionality in transposon silencing. These findings indicate that piRNA length specification, which is crucial for transposon repression, is conserved between species, but that the means of piRNA length specification differ between species.

## Results

### Daed and Gasz bind with each other via their respective eCC domains

According to the current model, the Daed–Gasz heterodimer tethers Armi bound to pre-Piwi-piRISC to mitochondria for Zuc-dependent piRNA processing.^12,29,30^ In this study, we first predicted the higher-order structure of the Daed–Gasz heterodimer using AlphaFold2^31,32^ (Figure 1A). This indicated that both Gasz and Daed have two α-helixes and that the coiled-coil structure composed of these four α-helixes is the center of the Daed–Gasz association (Figure 1B). Daed has SAM, coiled-coil (CC), and TM domains, and Gasz has ankyrin repeats (ARs), SAM, and TM domains.^12,29^ The predicted structures of Daed and Gasz showed that the second of the two α-helixes (Ala191– His235) of Daed roughly overlaps with the CC domain (Thr206–Ser234) and that the two α-helixes in Gasz (Phe345–Glu374 and Glu383–Ser412) are present between the SAM and TM domains (Figure 1C). The amino acid sequence-based prediction of the CC region^33^ also roughly overlaps with the tandem α-helix regions (Figure S1A), supporting the AlphaFold2 prediction. We propose that the two tandem α-helixes present in both Gasz and Daed are referred to as extended coiled-coil (eCC) domains (Figure 1C).

**Figure 1.**
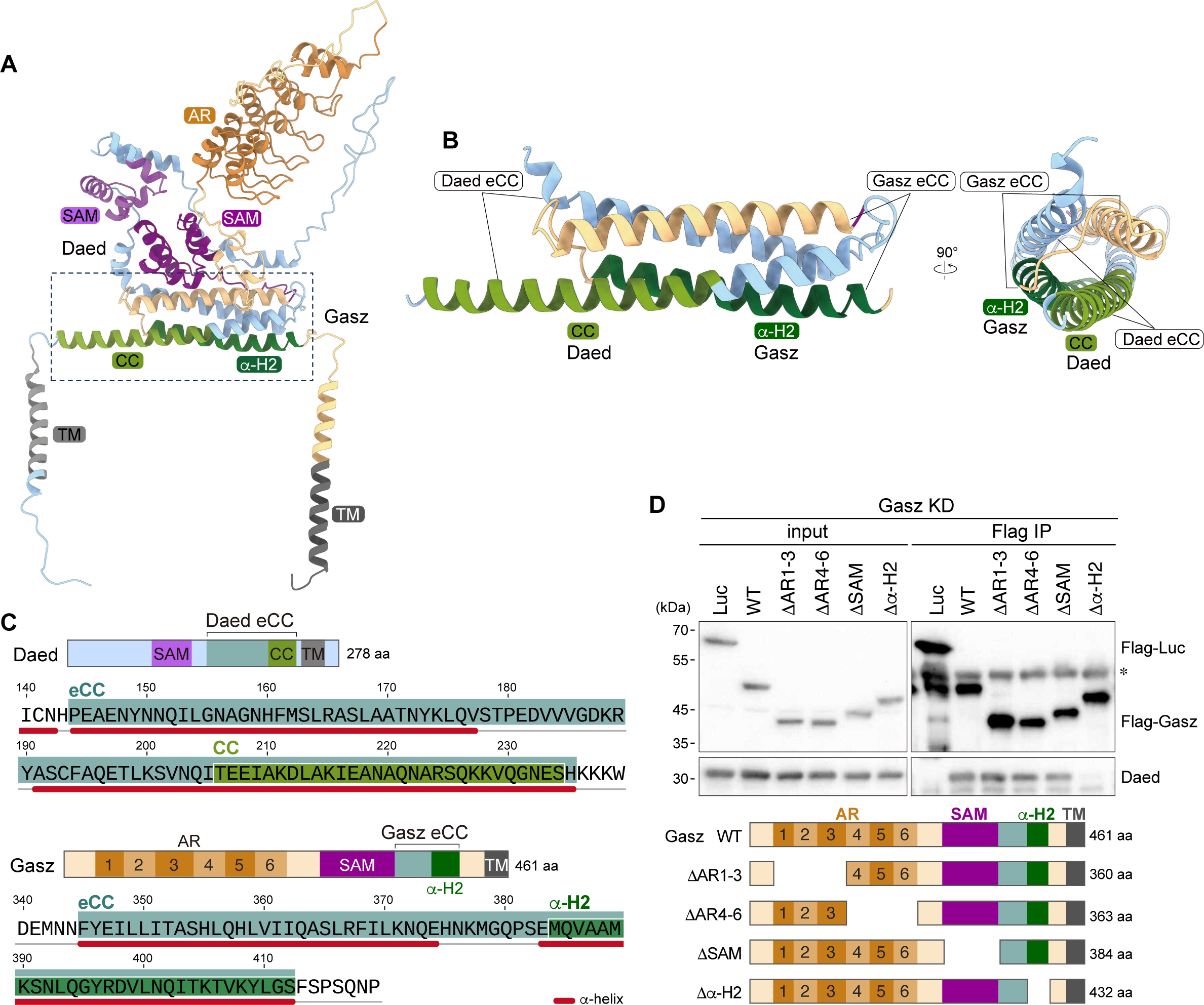
Daed and Gasz interact via their respective eCC domains. (**A**) Predicted structure of the Daed–Gasz heterodimer by AlphaFold2. (**B**) Magnification of the region indicated by the dotted line box in (**A**). The structure is shown from two directions. (**C**) Domain structures of Daed and Gasz. Peptide sequences of eCCs of Daed and Gasz are also shown. (**D**) Immunoprecipitation (Flag IP) and western blotting show that one of the Gasz mutants, Δα-H2, fails to bind to Daed in OSCs lacking endogenous Gasz. Flag-Luc: negative control. *: Heavy chains. The domain structures Gasz WT and deletion mutants are also shown.

The CC domain of Daed is the region that binds to Gasz.^29^ To identify the region of Gasz responsible for binding to Daed we expressed deletion mutants of Gasz in OSCs lacking endogenous Gasz and then performed immunoprecipitation assays. Three of the Gasz deletion mutants, ΔAR1-3, which lacks the first three ARs, ΔAR4-6, which lacks the second three ARs, and ΔSAM, bound to Daed as efficiently as wild-type (WT) Gasz did, but the other mutant, Δα-H2, lacking the second α-helix of eCC did not (Figures 1D and S1B). These results indicate that the respective eCC domains of Daed and Gasz are responsible for the Daed–Gasz association, consistent with the predicted structure.

### Daed stabilizes Gasz via protein–protein interaction

To verify whether Daed is required for piRNA biogenesis in cultured OSCs, we depleted intracellular Daed by RNA interference (RNAi) and conducted northern blotting. In control OSCs, both *idefix* piRNA (a representative nongenic piRNA) and *traffic jam* (*tj*) piRNA (a representative genic piRNA) were detected as expected (Figure S2A).^12^ However, in the absence of Daed, their piRNA precursors (pre-piRNAs) accumulated aberrantly, while mature piRNAs were barely detected (Figure S2A), as in OSCs lacking Gasz.^12^ This indicates that both Gasz and Daed are essential for piRNA maturation in OSCs.

Western blotting confirmed that RNAi was effective at depleting Daed and Gasz in OSCs (Figure 2A). We also found that, when Daed was depleted, levels of Gasz were also markedly reduced, but that levels of Daed were unchanged in the absence of Gasz (Figures 2A and 2B). Such trends were not observed for their mRNA levels (Figure 2C). These results indicate that Daed is required to stabilize Gasz at the protein level, but that Daed is not dependent on Gasz for its stabilization.

**Figure 2.**
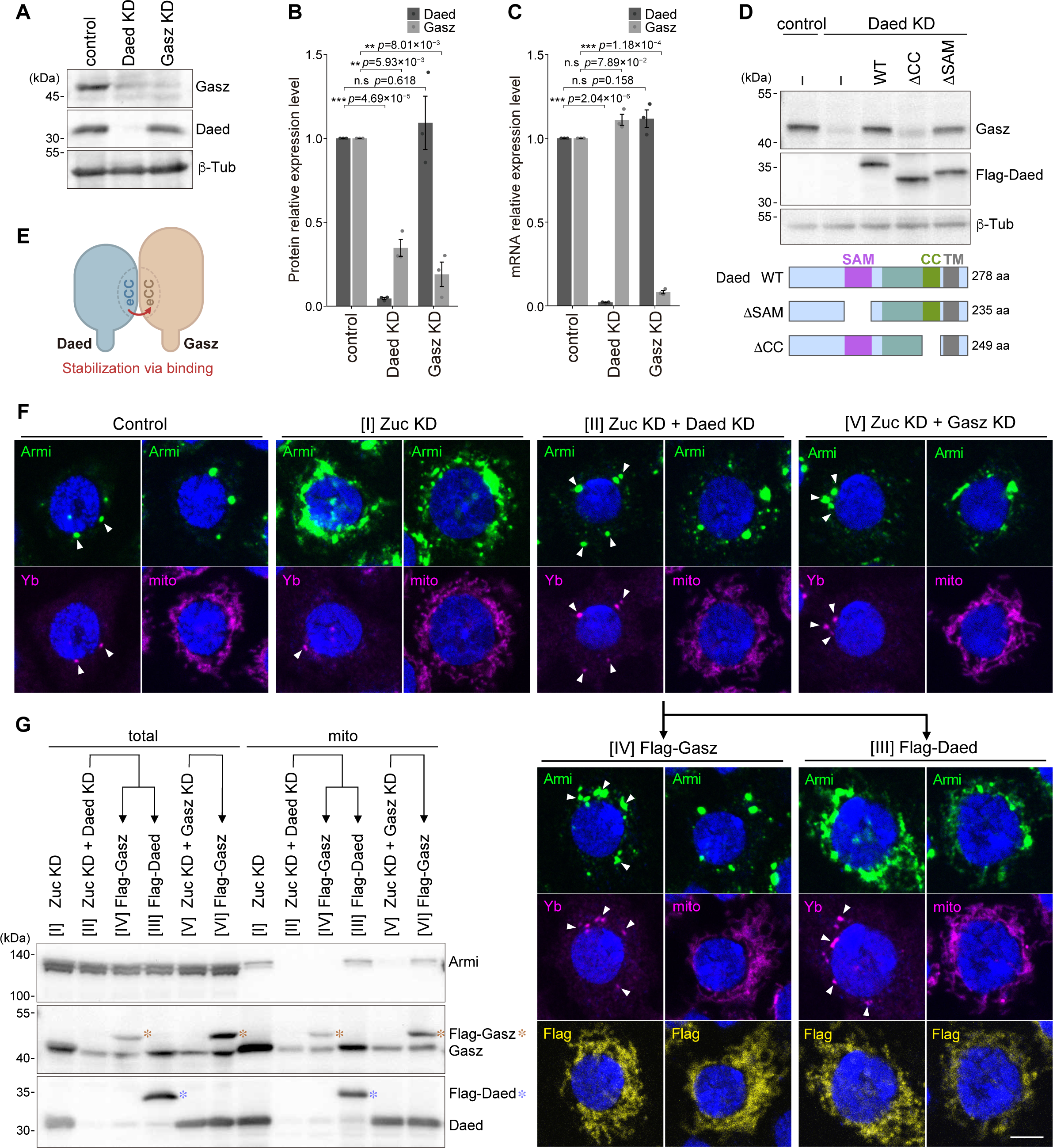
Daed stabilizes Gasz via interaction and both Daed and Gasz are essential for the mitochondrial tethering of Armi. (**A**) Western blotting shows the high efficiency of knockdown (KD) of Daed and Gasz in OSCs. Daed KD also destabilizes Gasz, but not *vice versa*. β-Tub: loading control. (**B**, **C**) The statistics of western blotting signals (n= 3; **A**) and the mRNA expression levels of *Daed* and *Gasz* upon Daed KD and Gasz KD (n=3). Signal intensities (**B**) or values (**C**) relative to control (EGFP KD) were normalized with those of β-Tub or *rp49*, respectively, and presented as mean values ± SE. **: *p* <0.01; ***: *p* <0.005; n.s.: *p* >0.05 [*t*-test (unpaired, two-sided)]. (**D**) Western blotting showing the levels of endogenous Gasz and Flag-Daed (WT and deletion mutants) in OSCs under each condition. β-Tub: loading control. The domain structures of Daed WT and mutants are shown. (**E**) Model of Gasz stabilization by Daed. (**F**) Subcellular localization of Armi (green) in OSCs under various conditions, [I] to [V]. Mitochondria (mito) and Yb bodies (Yb; white arrowheads) are shown in magenta. The signals of Flag-Gasz and Flag-Daed are shown in yellow. Images in the same vertical row are identical cells. Control: normal OSCs. DAPI (blue): nuclei. Scale bar 5 μm. (**G**) Western blotting shows the levels of endogenous Armi, Gasz, Daed, Flag-Gasz (orange asterisk), and Flag-Daed (blue asterisk) in total lysates (total) and mitochondrial fractions (mito) under conditions [I] to [VI]. For the efficiency of cell fractionation, see Figure S2C.

To investigate which domain of Daed is involved in Gasz stabilization, two deletion mutants of Daed, ΔSAM and ΔCC (part of the second α-helix of the eCC, see in Figure 1C), were expressed in OSCs depleted of endogenous Daed. Levels of Gasz were restored with expression of WT Daed and ΔSAM mutant Daed, but not with ΔCC mutant Daed (Figure 2D). A previous study showed that Daed binds to Gasz via the CC region^29^ and together these findings indicate that the Daed–Gasz interaction via their respective eCC domains is crucial for the stabilization of Gasz (Figure 2E). The Δα-H2 mutant of Gasz, which lacks the binding domain to Daed, was rather unstable in OSCs, but this was not because of mRNA instability (Figure S2B), supporting the above notion.

### Daed alone or Gasz alone is insufficient to tether Armi to mitochondria

Armi has at least two roles in piRNA biogenesis in OSCs; one is to transport nascent Piwi to Yb bodies to promote pre-Piwi-piRISC assembly, and the other is to translocate the complex from Yb bodies to mitochondria to promote Zuc-dependent piRNA processing.^12,34^ After this second role, Armi returns to the cytosol to repeat these functions, although immunofluorescence shows that Armi appears to be a constant resident of Yb bodies (Control, Figure 2F).^8,12^ In the absence of Zuc, Armi signals merged with mitochondrial signals {[I] Zuc knockdown (KD), Figure 2F}. This demonstrates dynamic movement of Armi in OSCs and indicates that Armi remains on the mitochondrial surface when loss of Zuc impairs piRNA maturation.^12^ Results from mitochondrial fractionation followed by western blotting were consistent with this statement (Figures 2G and S2C).

We then depleted Daed in Zuc-lacking OSCs, where both Daed and Gasz should be absent (Figure 2A). Under these circumstances, as expected, Armi was not retained on mitochondria and appeared in Yb bodies ([II] Zuc KD + Daed KD, Figure 2F). To restore the levels of both Daed and Gasz simultaneously, Flag-Daed was exogenously expressed in the OSCs and western blotting confirmed the recovery of both Daed and Gasz levels (Figures 2G and S2C). Immunofluorescence showed that Armi re-localized to mitochondria ([III] Flag-Daed, Figure 2F). However, when Gasz was forcibly expressed alone in the absence of Daed, Armi remained in Yb bodies ([IV] Flag-Gasz, Figure 2F). A similar result was obtained in OSCs expressing only Daed ([V] Zuc KD + Gasz KD, Figure 2F). These results indicate that Gasz alone or Daed alone is not sufficient to tether Armi to mitochondria. In other words, both Daed and Gasz are required to retain Armi on the mitochondrial surface for Zuc-dependent piRNA processing. Hereafter, the Daed–Gasz heterodimer is referred to as the Armi mitochondrial tethering complex (AmTEC). The behavior of Armi in OSCs under the conditions described above is summarized in Figure S2D.

### Daed, but not Gasz, binds to Armi via two distinct sites

We next predicted the higher-order structure of AmTEC bound with Armi (Figure 3A). Notably, no significant interaction between Gasz and Armi was observed, but Daed and Armi were bound at two distinct sites: The first site was through the first α-helix of eCC (Phe144–Val177) (Figure 1C), which interacted with the N-terminal region of Armi (Met1–Leu34) (Figures 3A, 3B, and S3A). The second interaction was between the SAM domain of Daed (Figure 1C) and the C-terminal helicase (i.e., UPF1-like family) domain of Armi (Figures 3A, 3C, and S3A).

**Figure 3.**
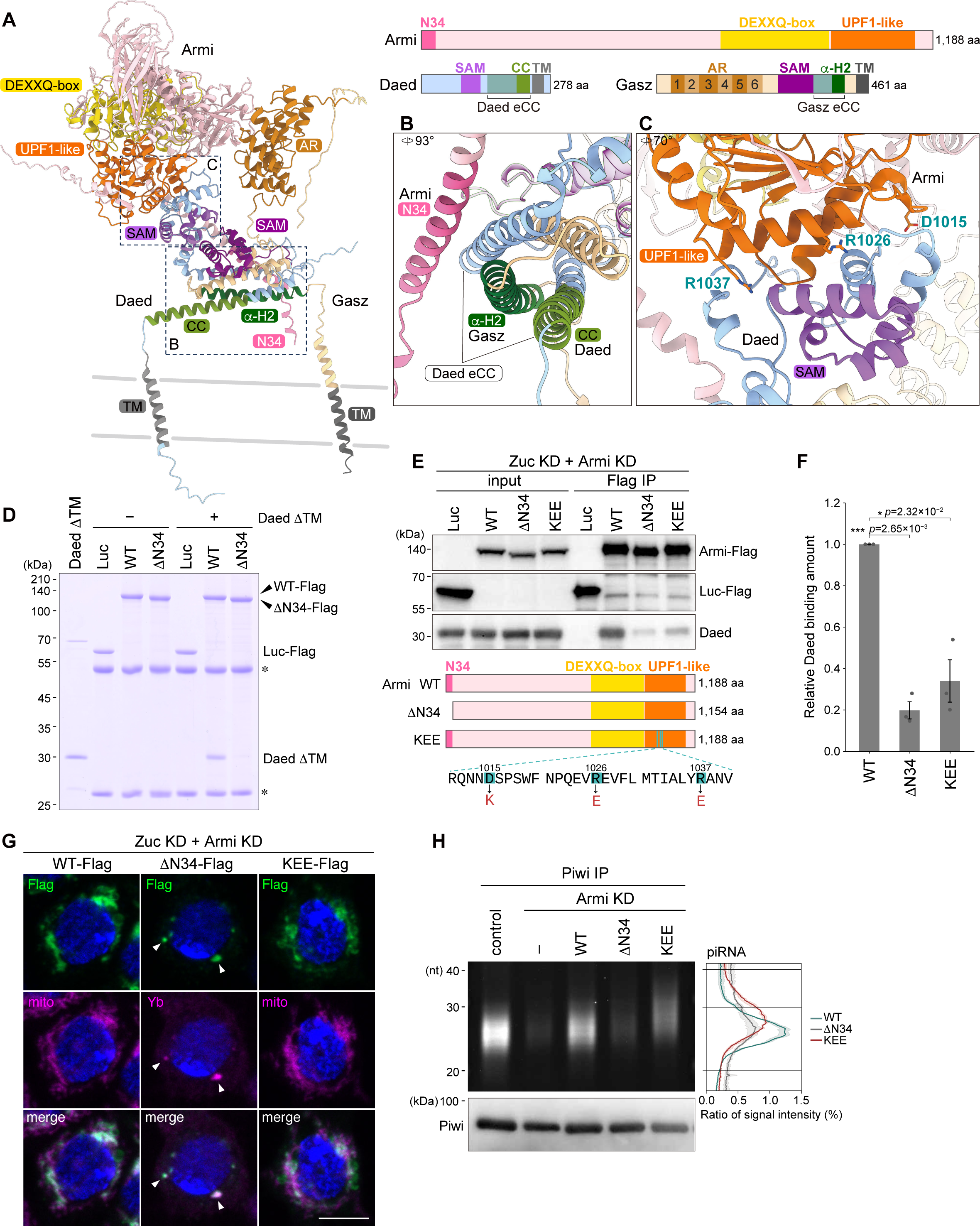
Daed interacts with Armi through two sites, each contributing differently to piRNA biogenesis. (**A**) The structure of the Armi–AmTEC complex predicted by AlphaFold2. Gray lines: virtual lipid bilayers. The domain structures of Armi, Daed, and Gasz are also shown. (**B**) Magnification of the “B” region in (**A**). The 34 N-terminal residues of Armi (N34) interacts with the first α-helix of Daed eCC. (**C**) Magnification of the “C” region in (**A**). The Armi UPF1-like helicase domain interacts with Daed SAM. Asp1015, Arg1026, and Arg1037 of Armi are shown in blue. (**D**) *In vitro* pull-down assays show Armi WT-Flag, but not Armi ΔN34-Flag, binding to Daed (Daed ΔTM-His). Luc-Flag: negative control. *: Heavy and light chains. (**E**) Immunoprecipitation (Flag IP) and western blotting show the amounts of endogenous Daed co-immunoprecipitated with Armi WT-Flag, ΔN34-Flag, and KEE-Flag. Luc-Flag: negative control. The domain structures of Armi WT and its mutants are shown below. Asp1015, Arg1026, and Arg1037 mutated in the KEE mutant are also shown. The peptide sequence of Armi is shown in Figure S3A. (**F**) The statistics of western blotting signals (n= 3; **E**). The signals of Daed were normalized with those of Armi and presented as mean values ± SE. *: *p* <0.05; ***: *p* <0.005 [*t*-test (unpaired, two-sided)]. (**G**) Subcellular localization of Armi WT, ΔN34, and KEE (green) in OSCs lacking endogenous Zuc (Zuc KD) and Armi (Armi KD). Mitochondria (mito; for WT and KEE) and Yb bodies (Yb, white arrowhead; for ΔN34) are shown in magenta. Images in the same vertical row are identical cells. DAPI (blue): nuclei. Scale bar: 5 μm. (**H**) Piwi-bound piRNAs obtained under each condition. Signal intensities (n=3) are normalized as a percentage of the total. The mean (line) and 95% confidence interval (shadow) are shown. The amount of Piwi in each sample is shown at the bottom.

To verify that the Armi–Daed interaction was via the Armi N-terminus (Figure 3B), recombinant WT Armi (WT-Flag) and its mutant lacking 34 N-terminal residues (ΔN34-Flag) were subjected to pull-down assays with Daed (Daed ΔTM) as prey. The Daed TM was removed because its inclusion interfered with protein preparation. Under conditions where a negative control, Luciferase (Luc), showed no binding to Daed ΔTM, Armi WT bound to Daed but Armi ΔN34 did not (Figure 3D). This confirms that Armi and Daed interact through the 34 N-terminal residues of Armi. We have previously shown that the Armi ΔN34 mutant (i.e., a shorter isoform of Armi) is unable to leave Yb bodies.^12^ The present analysis raises an alternative hypothesis; the Armi ΔN34 mutant may be able to leave Yb bodies but not be tethered to mitochondria because of the absence of Daed binding. This would explain why the shorter Armi isoform is inactive in piRNA biogenesis in OSCs.

To verify the Armi–Daed interaction via the Armi UPF1-like helicase domain (Figure 3C), three residues in the domain, Asp1015, Arg1026, and Arg1037 (D-R-R), which appear to be key to the binding, were mutated to Lys-Glu-Glu (K-E-E) and immunoprecipitation assays were conducted (Figure 3E). Quantification of the results indicated that the Armi KEE mutant bound to Daed with approximately 34% of the binding efficiency of WT Armi (Figure 3F). Meanwhile, the Armi ΔN34 mutant bound to Daed with approximately 20% of the binding efficiency of WT Armi. Immunofluorescence showed that the Armi ΔN34 mutant accumulated in Yb bodies more intensely than the KEE mutant in Arm/Zuc-lacking OSCs (Figure 3G). For binding to Daed, Armi may be more dependent on the N-terminal region than on the UPF1-like helicase domain.

A lack of Armi in OSCs severely disrupted piRNA production.^16,35^ This was almost fully rescued by the expression of WT Armi (Figure 3H). We next expressed the ΔN34 or KEE mutant of Armi as alternatives to the WT. The ΔN34 mutant failed to rescue the disruption to piRNA production, but the KEE mutant partially restored piRNA biogenesis (Figures 3H and S3B). This supports the notion that the N-terminal region of Armi is more important than the C-terminal helicase domain in piRNA biogenesis in OSCs. The strength of the Armi–Daed interaction (Figure 3E) appears to be positively correlated with the function of Armi.

### The two Armi–Daed associations differ in their functional contributions to piRNA biogenesis

piRNAs produced under conditions involving the Armi KEE mutant were notably increased in length by several bases (Figure 3H). We therefore investigated whether a similar phenomenon occurred under conditions involving the Daed ΔSAM mutant, given that the SAM domain is the binding partner of the UPF1-like helicase domain of Armi. Indeed, in cells expressing the Daed ΔSAM mutant as an alternative to endogenous Daed, Piwi-bound piRNAs became several bases longer and their levels were lower than those in cells expressing WT Daed, but higher compared with cells lacking endogenous Daed (Figures 4A and S4A).

**Figure 4.**
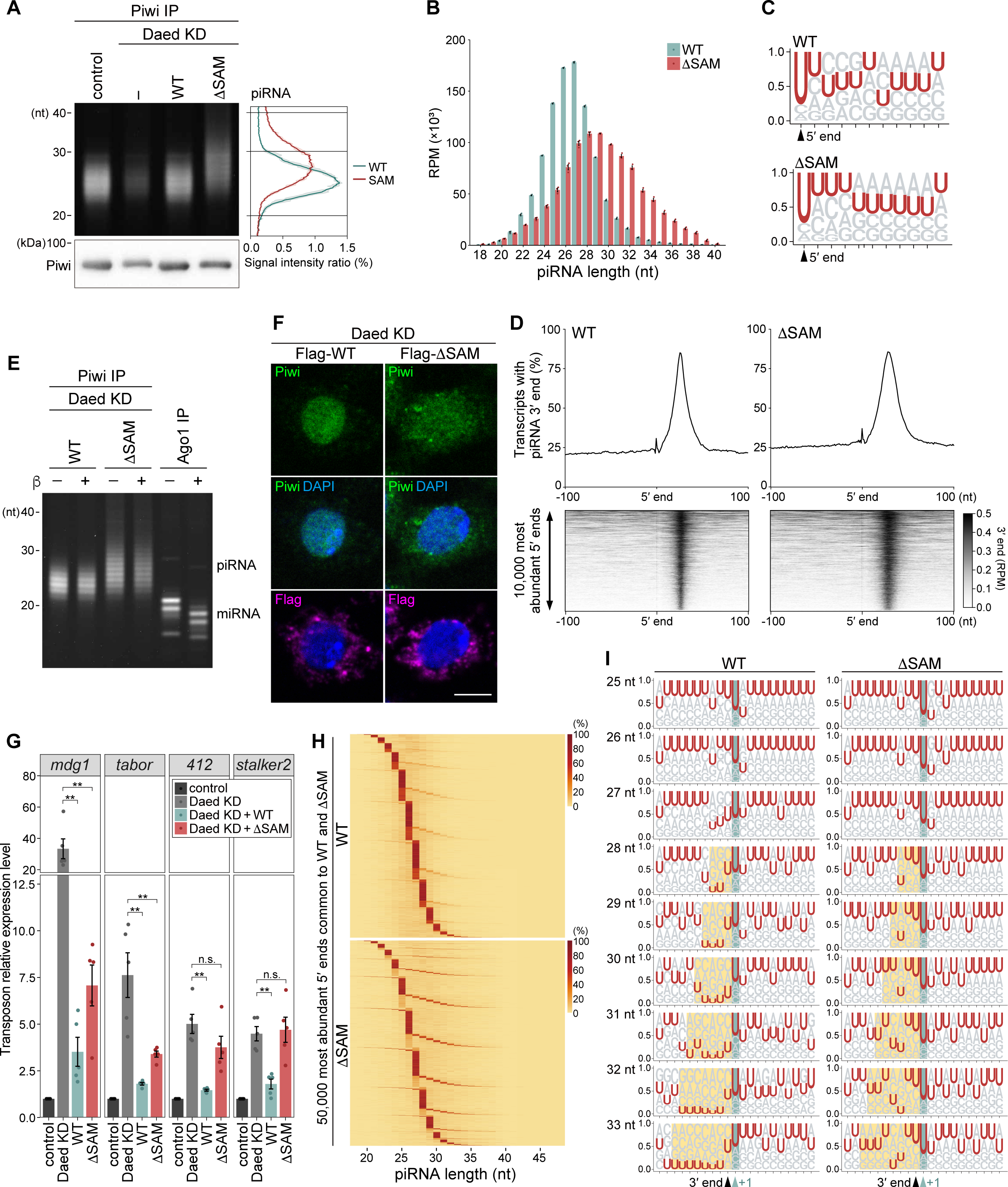
The loss of Armi–Daed interaction via Daed SAM obscures Zuc cleavage and impairs Piwi-piRISC function. (**A**) Piwi-bound piRNAs obtained under each condition. Signal intensities (n=3) are normalized as a percentage of the total. The mean (line) and 95% confidence interval (shadow) are shown. The amount of Piwi in each sample is shown at the bottom. (**B**) Length distribution of piRNAs (18–40 nt) in Daed WT (green) and ΔSAM (red) libraries (n= 3), presented as mean values ± SE. (**C**) Sequence logos representing 11 nt from the 5′ end of piRNAs in Daed WT and ΔSAM libraries. (**D**) Heatmaps showing 3′ end counts of piRNAs mapped around the 10,000 most abundant piRNA 5′ ends (lower) and its binary histograms (upper) in Daed WT (left) and ΔSAM (right). (**E**) β-elimination assays. Piwi-bound piRNAs in Daed KD OSCs expressing Daed WT and ΔSAM are shown. Ago1-bound miRNAs are used as positive controls. (**F**) Subcellular localization of endogenous Piwi (green) in Daed KD OSCs expressing Daed WT and ΔSAM (magenta). DAPI (blue): nuclei. Scale bar: 5 μm. (**G**) The expression levels of transposons (*mdg1*, *tabor*, *412*, and *stalker2*) in control OSCs (dark gray), Daed-lacking OSCs (Daed KD; light gray), Flag-Daed WT rescued (WT; green) or Flag-Daed ΔSAM rescued cells (ΔSAM; red) (n=5). Values relative to the control (EGFP KD) were normalized to *rp49* and presented as mean values ± SE **: *p* <0.01; n.s.: *p* >0.05 [Wilcoxon rank sum exact test (two-sided)]. (**H**) Heatmaps showing the frequency of the 3′ end positions (red) corresponding to each 5′ end of piRNAs in Daed WT (upper) and ΔSAM (lower) libraries. The 50,000 piRNA 5′ end positions common to both libraries and most abundant in the WT library were used for analysis. (**I**) Sequence logos. piRNAs were sorted by their lengths and aligned according to the 3′ end of the piRNAs (black arrowhead) and its immediately downstream nucleotide is highlighted in green. The U depletion found in 28– 33-nt piRNAs in the WT library and the equivalent regions in ΔSAM are highlighted in yellow.

To examine how the lack of binding between Daed SAM and Armi UPF1-like helicase domain actually affects piRNA biogenesis, we sequenced and analyzed Piwi-bound piRNAs in OSCs expressing Daed ΔSAM and compared the results with those from OSCs expressing WT Daed (Figures 4A and S4A). This length distribution of piRNAs was indeed shifted towards the longer fractions when they were produced in cells lacking the Daed SAM domain (Figure 4B). This trend was also observed when piRNAs were subdivided by target transposons (Figure S4B). In contrast, piRNA-mapped transposons were essentially unchanged (Figure S4C). This seems reasonable given that piRNAs are produced from *flam* in both cases. Furthermore, the 1U bias^5,36^ and piRNA phasing^17,18^ were similarly observed in both libraries (Figures 4C and 4D). The β-elimination experiments showed the 3′ ends of the piRNAs to be modified with 2′-*O*-methylation in both cases, unlike Argonaute 1 (Ago1)-bound miRNAs (Figure 4E).^37,38^ However, it should be noted that non-negligible amounts of Piwi immunofluorescence remained in the cytoplasm when Daed ΔSAM was expressed in OSCs lacking endogenous Daed (Figure 4F).

piRNA loading is essential for Piwi nuclear localization.^19,34^ Therefore, we next examined the piRNA binding status of cytoplasmic and nuclear Piwi. In contrast to nuclear Piwi, which was bound to piRNAs even in the long fraction, cytoplasmic Piwi was bound to few piRNAs (Figure S4D). This indicates that piRNAs produced under the ΔSAM conditions can localize Piwi to the nucleus as piRISC, regardless of length, but are insufficient to convert all Piwi to functional Piwi-piRISC. Consistent with this, transposons were not fully repressed under such conditions (Figures 4G and S4E). These results indicate that the Armi–Daed association via the UPF1-like helicase domain of Armi and the SAM domain of Daed is influential in Piwi-piRISC production and function, although less so than the other Armi–Daed association involving the 34 N-terminal residues of Armi and the eCC domain of Daed.

### The Armi–Daed interaction via Daed SAM optimizes Zuc endonucleolysis for piRNA-mediated transposon repression

To explore the nature of ΔSAM piRNA extension, we selected the 50,000 piRNA 5′ end positions common to the WT and ΔSAM and with high abundance in WT libraries (Figure S4F) and analyzed them further. Heatmaps depicted with their 5′ end positions aligned clearly indicate that ΔSAM piRNAs are more diverse in length. Many of them have some extra bases at the 3′ ends (Figure 4H). This is consistent with the length distribution shown in Figure 4A. There are also a few shorter piRNAs (Figure 4H). The relatively small number of shorter piRNAs may be because of protection by Piwi.

To further analyze the effects of Daed SAM loss on piRNAs, we extracted the 5′-end positions that dominantly produce 26- and 27-nucleotide (nt) piRNAs in the WT library, and then these positions were classified as “Unchanged” (51,833 positions) and “Elongated” (53,258 positions) according to the absence or presence of 3′-end elongation. piRNAs produced from both 5′-end groups maintained comparable phasing and +1U bias (i.e., 1U bias of piRNAs, real and virtual, immediately following piRNAs verified to bind to Piwi^17,18^) (Figure S4G), indicating that piRNA elongation caused by the loss of Daed SAM is caused by Zuc endonucleolysis.

We next sorted piRNAs by their length and aligned them according to the 3′ ends of piRNAs. This showed that atypically long piRNAs of around 28–33 nt in the WT library tended to have significantly low U bias near their 3′ ends (U depletion), just upstream of the 5′ ends of the immediately downstream piRNAs, as reported previously (yellow boxes in WT, Figure 4I).^18^ This was interpreted as showing that when Zuc cannot find U-rich regions in the piRNA precursors it waits to cleave the substrate until the U-poor region has passed, thereby producing atypically long piRNAs. This also explains why there were fewer 28–33-nt piRNAs in the WT library (Figure 4B). In contrast, there was less notable “U depletion” in 28–33-nt ΔSAM piRNAs (yellow boxes in ΔSAM, Figure 4I). In addition, there was a substantial number of 28–33-nt piRNAs in the ΔSAM library (Figure 4B). This indicates that when the Armi–Daed interaction is attenuated because of a lack of Armi–Daed association via Daed SAM (and the UPF1-like helicase domain of Armi), Zuc somehow produces these longer piRNAs more preferentially regardless of the U content in the precursor sequences while maintaining the +1U bias.

We simulated the effect of Daed ΔSAM on piRNA production. First, we hypothetically fragmented the sequence of the *flam* locus (21,629,891 to 21,792,731 on the X chromosome; T was converted to U) into 163 piRNA precursors, starting at U and into sizes of 1,000 nt or close to that size. In parallel, the data shown in Figure 4B were fitted to normal distributions. Then we simulated Zuc-dependent piRNA production from their 5′ ends by assuming that Zuc cleavage occurs before U, produces 18–40-nt piRNAs, and follows the normal distributions for WT and ΔSAM. Sequence analysis of the obtained piRNAs (25–34 nt) indicated that U depletion was observed in simulations fitting the WT distribution, starting from 29 nt, but less so in simulations fitting the ΔSAM distribution (Figure S4H). This suggests that Zuc has the potential to produce Piwi-piRISC without following the U-depletion rule.

To investigate potential causes of Zuc-dependent precursor cleavage changes in ΔSAM, molecular dynamics simulations were performed using the Armi–AmTEC complex model (Figure 3A) and its Daed ΔSAM version. The distance between the center of mass (CoM) of Armi helicase domain (Ser697–Cys1,148 in Figure S3A) and that of the AmTEC was measured (n=5) and the average value was calculated at all time points along each trajectory (Figures S5A and S5B). This analysis shows that there is little difference in the initial Armi–AmTEC CoM distances between Daed WT and ΔSAM, but as the simulation time progresses, the difference tends to increase for the ΔSAM version (Figure S5C). The statistical analysis also shows that the distribution of CoM distances at all time points for all trajectories shifts to larger values for ΔSAM (Figures S5D). These results suggest that Daed ΔSAM tethers Armi more precariously than Daed WT. Based on these results, we hypothesize that “proper” binding of Armi and Daed via the Daed SAM (and the UPF1-like helicase domain of Armi) is critical for maintaining the distance between Armi and the scaffold complex, the loss of which would obscure the location of piRNA precursor cleavage by Zuc, negatively influencing Piwi-piRISC production.

We also sequenced and analyzed Piwi-bound piRNAs in OSCs expressing the Armi KEE mutant or WT Armi (Figures 3H and S3B). We found that piRNAs in the Armi KEE library were also several bases longer than the Armi WT piRNAs and retained 1U bias (Figures S6A and S6B). Heatmaps showed that the KEE piRNAs also exhibited characteristics of 3′ end elongation (Figure S6C). These results further support our notion that the SAM domain of Daed and the UPF1-like helicase domain of Armi are the binding partners and that the Armi–Daed interaction is indispensable to optimize Piwi-piRISC maturation by Zuc and piRISC function in OSCs (Figure S6D).

### Daed is Diptera-specific and controls Zuc cleavage in a Nibbler-independent manner

Armi and Gasz are widely conserved among species from invertebrates to vertebrates, but Daed is unique to the Diptera order, which is composed of Brachycera suborder, which includes *Drosophila* and other fly species, and Nematocera suborder which includes mosquitoes (Figures 5A, S7A, and S7B). In contrast, other species not belonging Diptera order do not appear to have Daed orthologs.^39^

**Figure 5.**
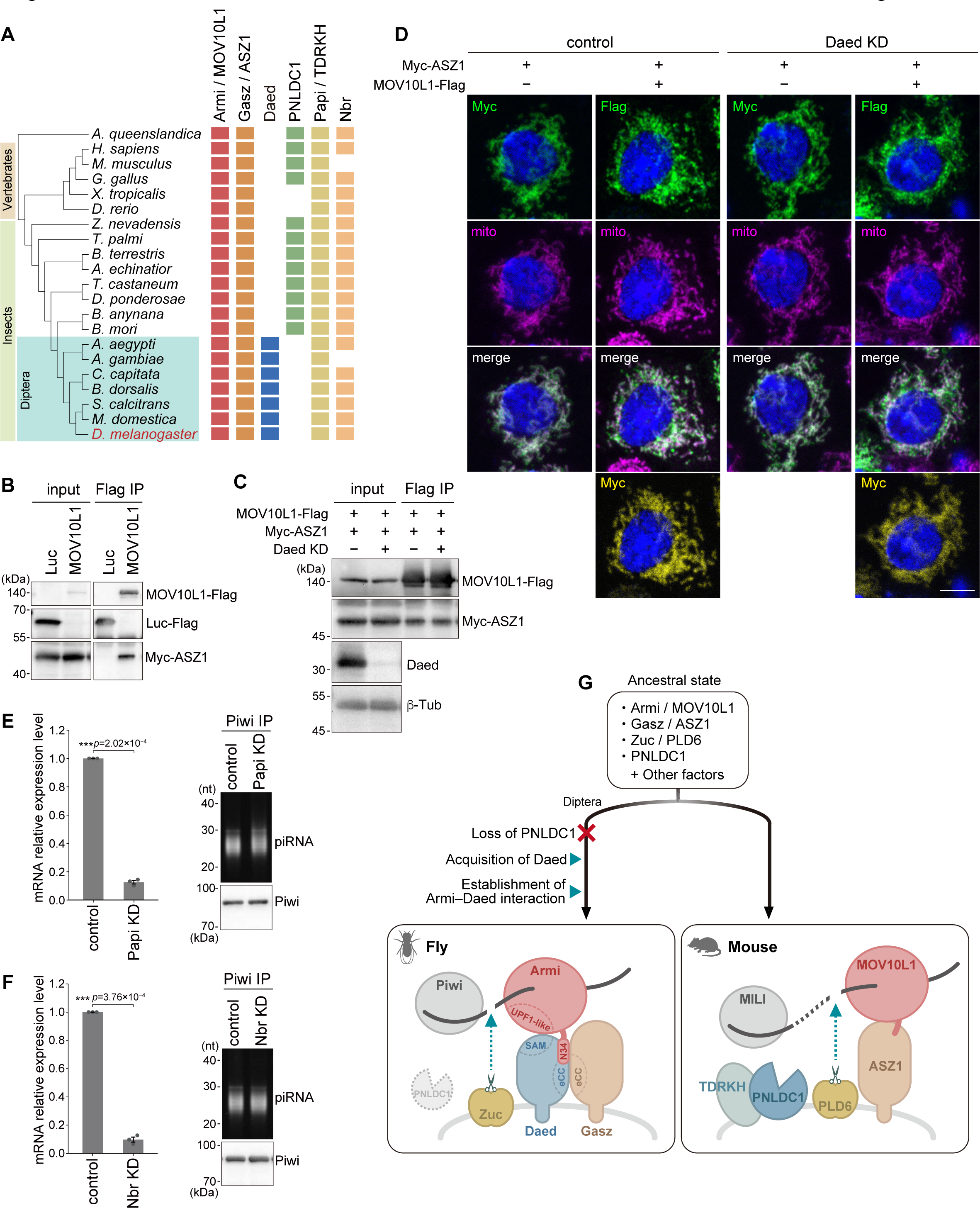
Daed is Diptera-specific and controls Zuc cleavage in a Nbr-independent manner. (**A**) Protein conservation state of Armi, Gasz, Daed, PNLDC1, Papi, and Nbr in *D. melanogaster* and other species. (**B**, **C**) Immunoprecipitation (Flag IP) and western blotting show Myc-ASZ1 binds to MOV10L1 in normal (**B**) and Daed-depleted OSCs (**C**). Luc-Flag: negative control. (**D**) Subcellular localization of Myc-ASZ1 alone or MOV10L1-Flag (green) with Myc-ASZ1 (yellow) in normal (left two row) and Daed-depleted OSCs (right two row). Mitochondria (mito) are shown in magenta. Images in the same vertical row are identical cells. DAPI (blue): nuclei. Scale bar: 5 μm. (**E**, **F**) The efficiency of Papi KD (upper) and Nbr KD (lower) (n=3). Piwi-bound piRNAs in control and Papi-KD (upper) and Nbr-KD OSCs (lower). The amount of Piwi in each sample is also shown. (**G**) Model proposed in this study. Diptera species acquired Daed instead of PNLDC1/PARN during evolution; Daed and Gasz form AmTEC, which stably tethers Armi to the mitochondrial surface via two sites. This allows Zuc to define the length of piRNAs without relying on exonucleases such as PNLDC1/PARN or Nbr. In contrast, in mice, MOV10L1/Armi is tethered solely by ASZ1/Gasz. In this case, Zuc cleavage does not specify piRNA length; therefore, trimming by exonucleases such as PNLDC1/PARN is required after Zuc processing. Blue arrowhead: Zuc cleavage.

In mice, primary piRNA biogenesis occurs in the inter-mitochondrial cement, where Armi homolog, MOV10L1, Gasz homolog, ASZ1, Zuc homolog, PLD6, and TDRD proteins play a role in piRNA production: MOV10L1 binds to PIWI proteins (MILI, MIWI and MIWI2) and piRNA precursors and contributes to piRNA biogenesis through its helicase activity.^40,41^ ASZ1 is essential for the production of MILI-bound piRNAs in mice^42^ and is thought to tether MOV10L1 to mitochondria, together with TDRD proteins, such as TDRD1, to promote MIWI-piRISC production.^40,41^

We expressed both MOV10L1 and ASZ1 in OSCs and performed immunoprecipitation. The MOV10L1–ASZ1 interaction was detected to a similar extent in both normal OSCs and OSCs lacking intracellular Daed (Figures 5B and 5C). Immunofluorescence confirmed this; the expression of ASZ1 was sufficient for the mitochondrial localization of MOV10L1, independent of the presence of intracellular Daed (Figure 5D). This indicates that the MOV10L1–ASZ1 interaction and MOV10L1 tethering is Daed-independent, while the Armi–Gasz interaction and Armi tethering in *Drosophila* required Daed. These results also indicate that the Armi–Gasz association, at least in OSCs, does not require other factors present in mouse testis. ASZ1 was stable in OSCs lacking endogenous Daed (Figure 5C), unlike Gasz (Figure 2A), indicating that the molecular system that actively degrades Gasz in the absence of Daed does not act on ASZ1.

Why did *Drosophila* acquire Daed during evolution? In mice, poly(A)-specific ribonuclease-like domain containing 1 (PNLDC1), a homolog of poly(A)-specific ribonuclease (PARN), trims the 3′ ends of piRNAs and piRNAs are elongated after PNLDC1 loss of function.^43,44^ Mutation in PNLDC1 reduces LINE-1-targeting piRNAs and mislocalizes MIWI2 to the cytoplasm, leading to LINE-1 derepression. This was nearly equal to what we observed in this study with the *Drosophila* Daed ΔSAM mutant. However, PNLDC1 is not conserved in the majority of Diptera order species.^45^ Papi is the *Drosophila* ortholog of TDRKH, which, in mice, acts as a scaffold for PNLDC1 and whose functional inhibition causes piRNA elongation.^46,47^ However, consistent with previous studies using Papi mutant flies,^45^ its depletion had little effect on piRNA length in OSCs (Figure 5E).

The 3′-to-5′ exonuclease, Nibbler (Nbr), trims both miRNAs and piRNAs in *Drosophila* and mosquitoes,^45,48–50^ but the pre-piRNAs that undergo trimming are minor in mosquitoes, and the loss of Nbr barely affected the length of Piwi-bound piRNAs in the *Drosophila* ovary.^45^ Our study confirmed this; namely, Nbr depletion in OSCs had little effect on the piRNA length (Figure 5F). This indicates that Nbr may function in the trimming of miRNAs, but not piRNAs, in OSCs. The lack of a marked increase in +1U signal in piRNAs elongated in the Daed ΔSAM mutant (Figures 4D and S4G) also supports this view.

In contrast, Armi D-R-R, which was essential for the regulation of Zuc cleavage in *Drosophila*, is highly conserved in the fly species that lack PNLDC1/PARN and have Daed (Figure S8). These results indicate that *Drosophila,* and its closely related species, may have acquired Daed after losing PNLDC1/PARN during evolution, and that they may also have acquired the ability to produce mature piRNAs without further trimming by exonucleases by allowing Armi to bind directly to Daed (Figure 5G).

## Discussion

This study revealed that Daed and Gasz bind to each other via their eCC domains, both of which are composed of two α-helixes that assemble into a stable helical barrel consisting of all four of these α-helixes. When Daed lacks the second α-helix in the eCC domain, the Daed–Gasz interaction is disrupted and Gasz becomes destabilized, illustrating the importance of Daed in Gasz stabilization (Figure 2E). This also shows that the intracellular level of Gasz is strictly regulated by the intracellular level of Daed, indicating that OSCs do not tolerate the presence of Gasz alone. In contrast, OSCs do tolerate the presence of Daed alone. However, Daed, in the absence of Gasz, failed to anchor pre-Piwi-piRISC to mitochondria via Armi. Gasz is therefore complicit in the role of Daed, although the contribution of Gasz to the direct binding of Daed to Armi appears to be minimal.

Lack of binding through the Daed SAM did not completely abolish the Armi– Daed association but resulted in aberrant Zuc function (Figure S6D). Under such conditions, Piwi-bound piRNAs were atypically long because of extra bases at the 3′ ends, and there was no notable U depletion, a typical feature of piRNAs in normal OSCs. In normal OSCs, atypically long piRNAs are produced only when piRNAs of typical sizes cannot maintain the +1U bias. In contrast, in OSCs expressing Daed ΔSAM, Zuc shows “relaxed selectivity” in terms of nucleotides and length and tends to produce longer piRNAs, even under conditions where it can produce shorter (i.e., normal size) piRNAs with the +1U bias, albeit in relatively small quantities. This could be because of the timing of Zuc RNA cleavage was shifted slightly downstream of the precursor while the preference for cleavage before U was conserved. Recombinant Zuc did not show U selectivity,^15^ indicating that, *in vivo*, Zuc may rely on a cofactor to confer U selectivity, and that this cofactor may not require the Armi–Daed association to execute its function.

Interestingly, Piwi could accept those longer piRNAs produced under the ΔSAM conditions and localized to the nucleus as Piwi-piRISC. At the same time, however, nonnegligible amounts of Piwi remained in the cytoplasm because of the lack of piRNA loading, which resulted in transposon derepression. This means that the ability of Daed to interact with Armi via the SAM domain, in addition to the binding via the first α-helix in the eCC domain, is essential not only to define the positions of Zuc cleavage, but also to produce enough Piwi-piRISC for transposon repression in OSCs. It is worth noting that no such regulatory factor has been identified to date.

Armi is unique in that it plays multiple roles in piRNA biogenesis. First, Armi delivers unloaded Piwi to Yb bodies, facilitating the assembly of pre-Piwi-piRISC in these organelles. In Yb bodies, Armi binds to pre-Piwi-piRISC and translocates the complex to mitochondria. There, Armi anchors pre-Piwi-piRISC to the mitochondria by binding to Daed. This allows Zuc to process pre-Piwi-piRISC into mature Piwi-piRISC. Armi is therefore a factor that always accompanies Piwi and encourages it along the pathway to develop into Piwi-piRISC.

Armi is an RNA helicase protein, and its RNA helicase activity is required to avoid stochastic interactions with abundant mRNAs.^51^ In contrast, up to this point, the RNA helicase activity of Armi does not seem to be required for the piRNA precursors, rather its activity would be inhibitory. However, in the absence of this activity, especially for phasing, the sequential generation of Piwi-piRISC from a single piRNA precursor would not occur. It is possible that an inhibitor of the RNA helicase activity of Armi is present in OSCs and functions until phasing, but, upon binding to Armi, Daed would lose its function allowing phasing to occur.

Animals that reproduce sexually retain the highly conserved piRNA mechanism and use it to maintain their fertility by repressing transposons. However, the evolutionary rate of many piRNA factors, including Armi, is rapid,^52,53^ suggesting that these species have adapted the machinery to their specific endogenous and exogenous environments. Coevolution of piRNAs and transposons through host–pathogen arms races has been proposed as a background to this process.^54^ Although flies and mosquitoes belong to the same Diptera order, the piRNA pathway factors are highly diverse.^55^ On the other hand, our observation that Daed is conserved among Diptera species may suggest the importance of this factor in Dipteran piRNA biogenesis pathway where PNLDC1 is lost.

## Author contributions

YK and MN performed biochemical experiments. YK, SH, and HY performed bioinformatic analyses, computational structural prediction, simulation, and calculation. MCS supervised and discussed the work and wrote the manuscript with the other authors.

## Acknowledgments

We thank Y. Namba, Y.W. Iwasaki, H. Nishimasu, and S. Kuraku for useful discussions, K. Kudo, T. Yasuda, and Y. Fukushima for technical assistance with the molecular dynamics simulation, and T. Fujisawa, T. Kinoshita, and H. Ishizu for technical assistance with the biochemical analysis. This study was supported by MEXT KAKENHI Grant Numbers JP19H05466 (to MCS), JP22K15056 (to SH), JP23K14141 (to HY), and JP22KJ1122 (to YK). Some computations were performed on the NIG supercomputer at ROIS National Institute of Genetics.

## STAR★Methods

### Key resources table

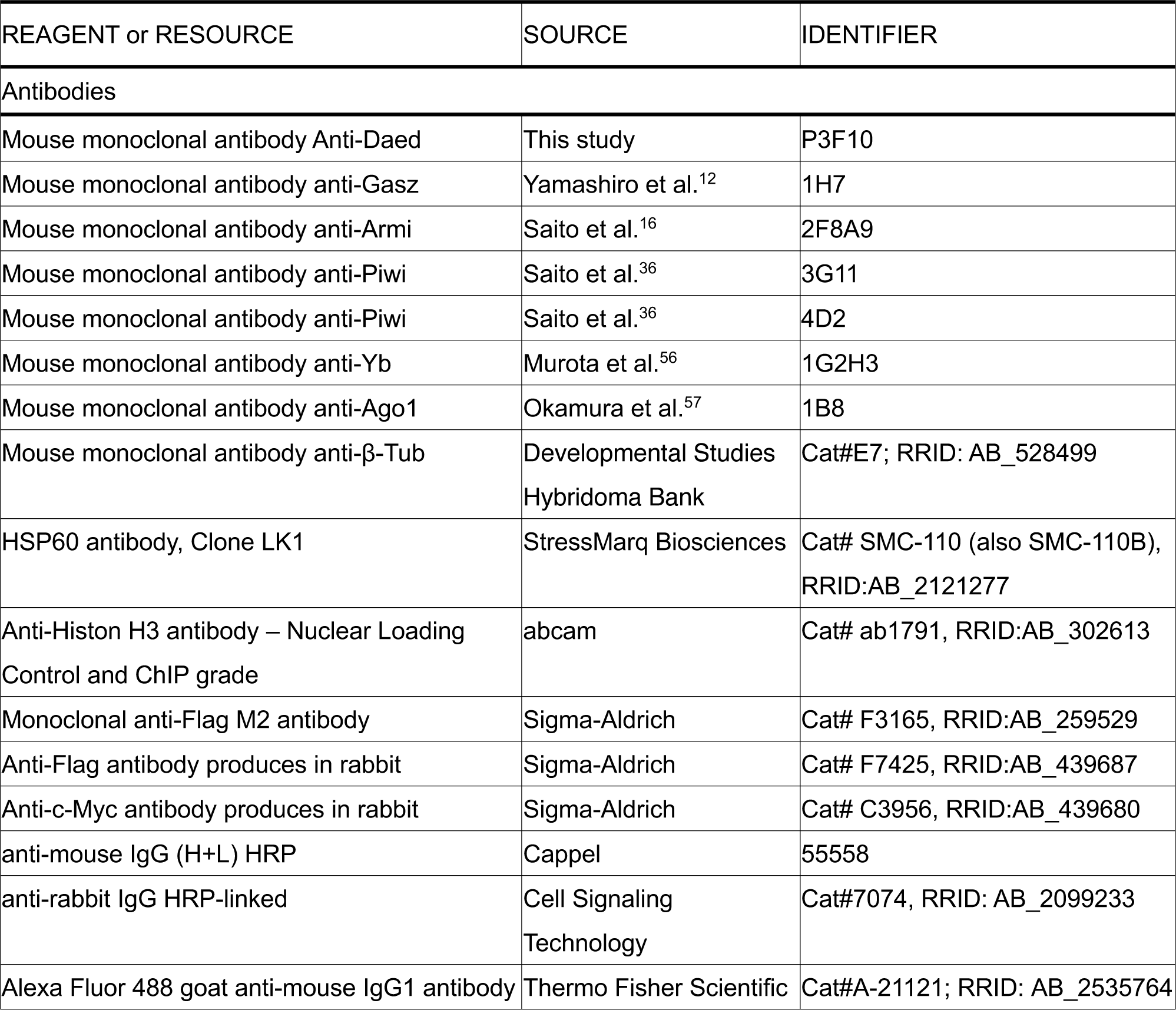
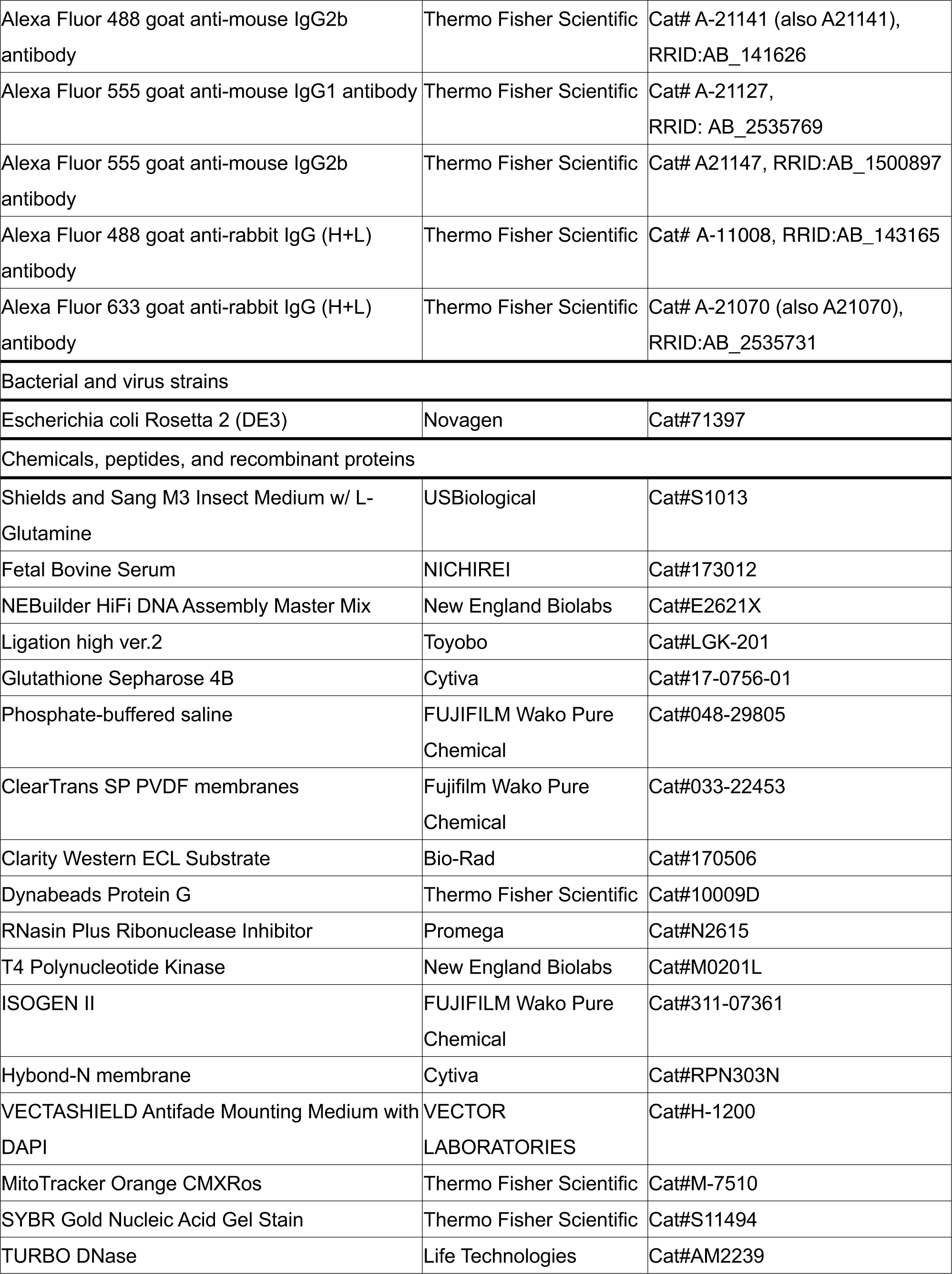
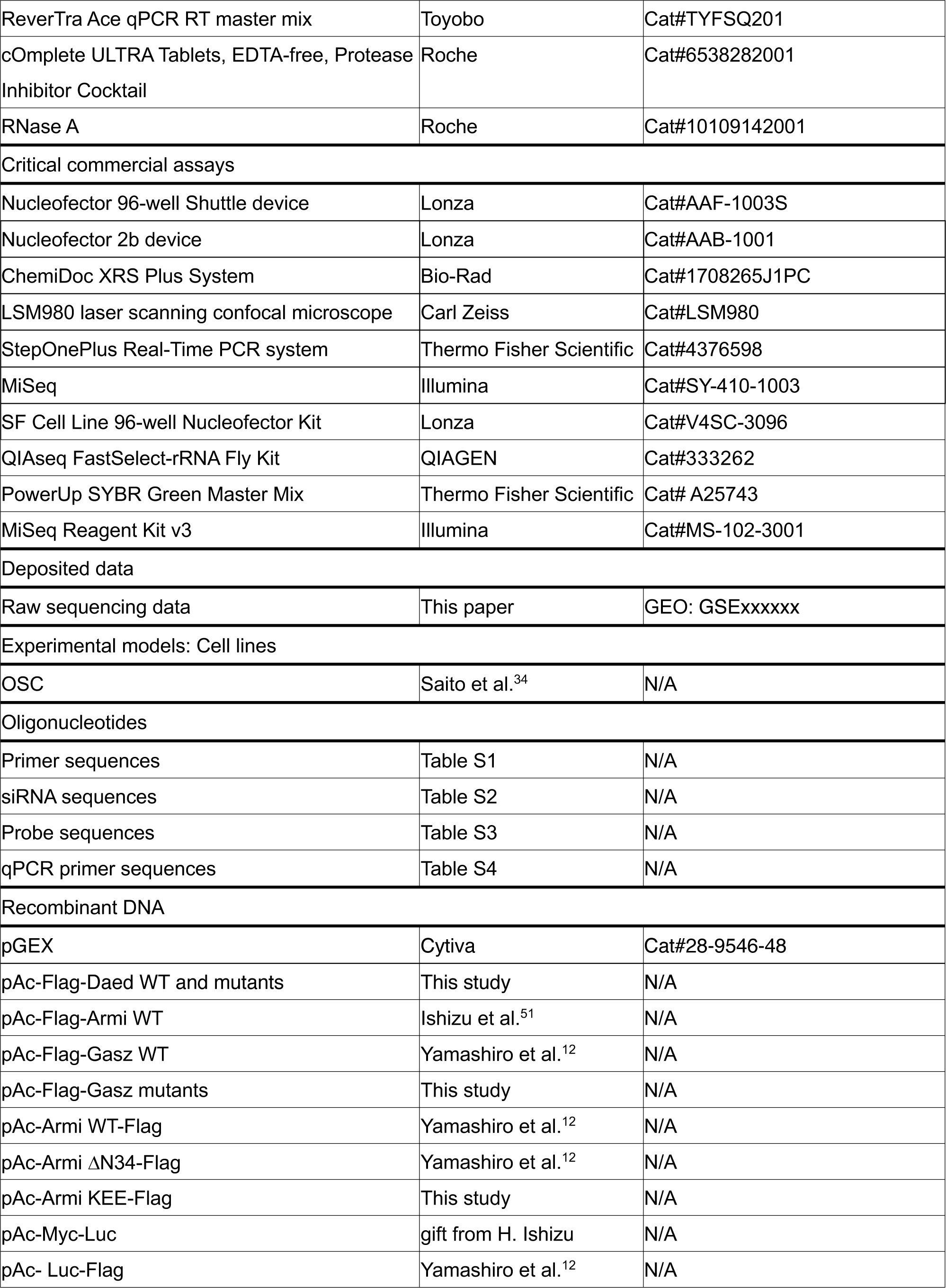
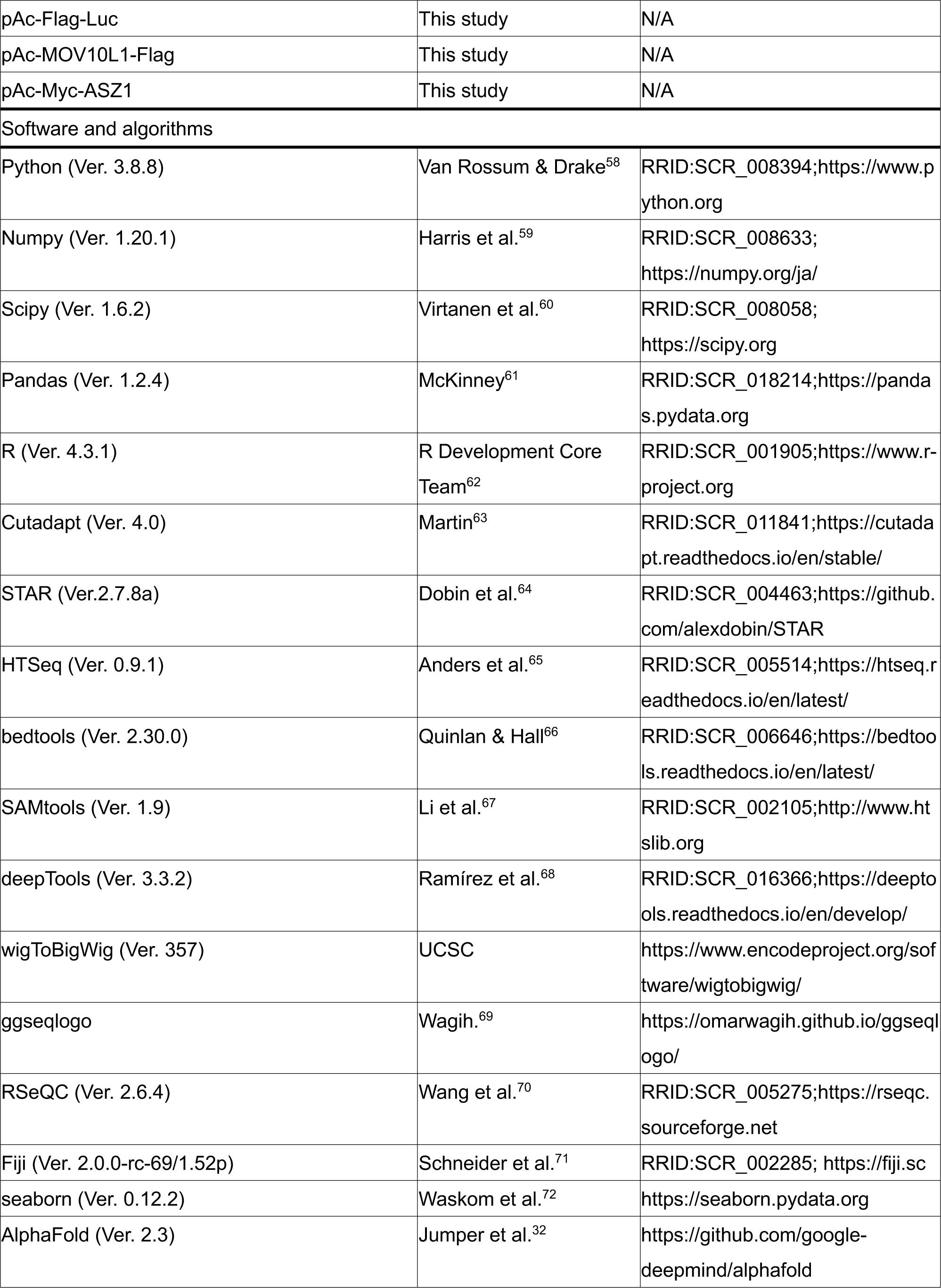
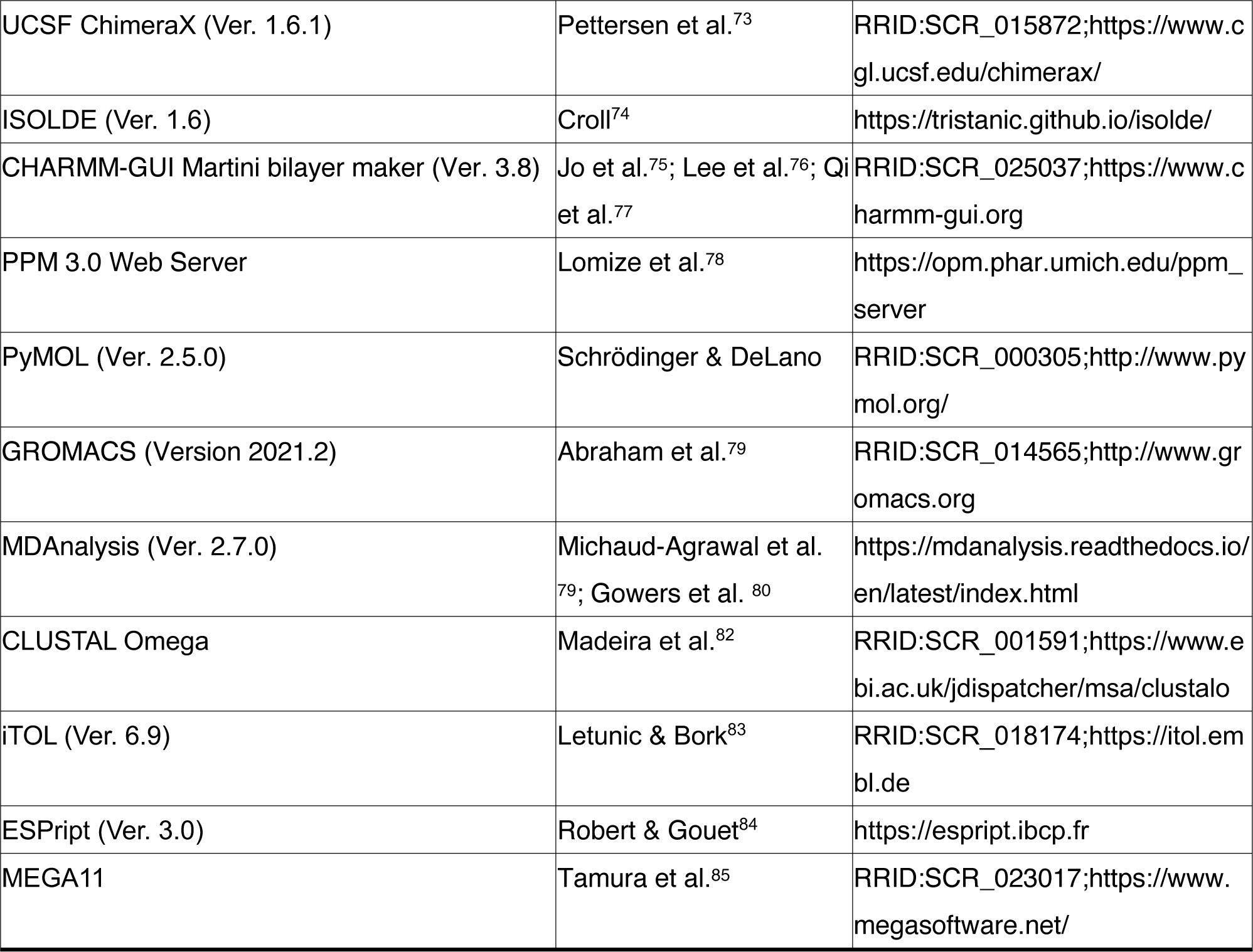

## Resource availability

### Lead contact

Further information and requests for resources and reagents should be directed to and will be fulfilled by the lead contact, Mikiko C. Siomi.

### Materials availability

Plasmids and antibodies generated in this study will be shared by the lead contact upon request.

### Data and code availability

The data supporting the findings of this study are available from the corresponding authors upon reasonable request. The datasets produced in this study are available in the following databases: RNA-Seq data: Gene Expression Omnibus GSExxxxxx (https://www.ncbi.nlm.nih.gov/geo/query/acc.cgi?acc=GSExxxxxx).

### Experimental model and study participant details

OSCs were cultured at 26°C in Shields and Sang M3 medium (USBiological) supplemented with 10% fetal bovine serum (FBS, NICHIREI), 0.6 mg/mL glutathione, 10 mU/mL insulin, and 10% fly extract.^8^

## Method details

### Plasmid construction

The Daed coding region was amplified from total OSC RNAs by RT-PCR and inserted into pAc-Flag-Armi WT^51^ by substituting the Armi coding region using NEBuilder HiFi DNA Assembly Master Mix (New England Biolabs). RNAi-resistant Daed WT and its ΔSAM and ΔCC mutants were generated by inverse PCR, according to the domain structures described in a previous study.^29^ To construct the expression vector for GST-Daed ΔTM-His, corresponding coding regions of Daed were cloned into pGEX, after which a His-Tag was added to the C-terminal by inverse PCR. RNAi-resistant Flag-Gasz mutants (ΔAR1-3, ΔAR4-6, ΔSAM, and Δα-H2) were generated by inverse PCR using RNAi-resistant Flag-Gasz WT vector^12^ as a template. Vectors for RNAi-resistant Armi WT-Flag and its ΔN34 mutant were previously constructed.^12^ The Armi KEE-Flag mutant was produced by inverse PCR using RNAi-resistant Armi WT-Flag vector as a template. The Flag-Luc expression vector was generated from pAc-Myc-Luc (a gift from H. Ishizu) by substituting Myc with 3×Flag. The MOV10L1 and ASZ1 coding region was amplified from total mouse ovary RNAs by RT-PCR and inserted into pAc-Armi WT-Flag and Myc-Daed WT by substituting original coding regions. The primers used are shown in Table S1.

### RNAi and transfection

To initiate RNAi, 5.0 × 10^6^ OSCs were mixed with 200 pmol of siRNAs in 20 μL of Solution SF of the Cell Line Nucleofector Kit SF (Lonza Bioscience) and electroporation was performed using a Nucleofector 96-well Shuttle device (Lonza Bioscience). For plasmid transfection or plasmid/siRNA co-transfection, 1.5 × 10^7^ OSCs were mixed with 1.5–10 μg of plasmids with or without 600 pmol of siRNAs in 100 μL of transfection buffer [180 mM sodium phosphate buffer for Church and Gilbert hybridization (pH 7.2), 5 mM KCl, 15 mM MgCl_2_, and 50 mM D-mannitol] modified from a previous report,^86^ and electroporation was performed by a Nucleofector 2b device (Lonza Bioscience). Cells were transfected with siRNAs twice with a 48-h interval. For rescue assays, endogenous genes were depleted by RNAi and plasmids and siRNAs were introduced into the cells after 48 h of culture. Cells were harvested after another 48 h of culture. For immunofluorescence in Figure 2F, 500 pmol of siRNAs were used for RNAi and plasmid/siRNA co-transfection, and cells were harvested after another 72 h of culture. Sequences of siRNAs are indicated in Table S2.

### Production of anti-Daed monoclonal antibody

GST-Daed (Met1–Asp197) was expressed in *E. coli* (Rosetta-DE3) and the inclusion bodies were used for mouse immunization. Production and selection of hybridomas that produce anti-Daed monoclonal antibodies were performed as described previously.^87^

### Mitochondrial fractionation

Mitochondria were fractionated as previously reported.^12^ Briefly, 1.2 × 10^8^ OSCs were suspended in 500 μL of buffer A [30 mM Tris-HCl (pH 7.3), 225 mM D-mannitol, 75 mM sucrose, and 0.05 mM EGTA] and the cell suspension was passed 30 times through a 30-gauge needle attached to a 1-mL syringe. The lysates were centrifuged at 600 *g* for 5 min and the supernatants were collected, and repeated twice. The lysates were then centrifuged at 7,000 *g* for 10 min and the supernatants were collected as the cytoplasmic fraction. This step was also repeated twice. The pellets were resuspended in 500 μL of buffer B [30 mM Tris-HCl (pH 7.3), 225 mM D-mannitol, and 75 mM sucrose] and centrifuged at 10,000 *g* for 10 min. The supernatant was used as the mitochondrial fraction.

### Nuclear fractionation

Nuclear fractionation was performed as previously reported with some modifications. In brief, cells were suspended in hypotonic buffer [10 mM HEPES-KOH (pH 7.3), 10 mM KCl, 1.5 mM MgCl_2_, 0.5 mM dithiothreitol (DTT), 2 μg/mL leupeptin, 2 μg/mL pepstatin A, and 0.5% aprotinin] by pipetting six times and passed 3 times through a 25-gauge needle attached to a 1-mL syringe. The lysates were centrifuged at 400 *g* for 10 min to collect nuclei. The supernatants were collected as the cytoplasmic fraction, then the concentration of KCl was adjusted to 200 mM and centrifuged at 20,000 *g* for 20 min before used for Piwi-piRISC isolation (see below). The nuclear fraction was washed with hypotonic buffer and resuspended in chromatin co-immunoprecipitation buffer [50 mM HEPES-KOH (pH 7.3), 200 mM KCl, 1 mM EDTA, 1% Triton X-100, 0.1% Na deoxycholate, 2 μg/mL leupeptin, 2 μg/mL pepstatin A, and 0.5% aprotinin]. The nuclear fraction was sonicated and centrifuged at 20,000 *g* for 20 min. The supernatant was collected as nuclear extract fraction for Piwi-piRISC isolation.

### Western blotting

Western blotting was performed essentially as described previously.^88^ Proteins separated on SDS polyacrylamide gels were transferred to ClearTrans SP PVDF membranes (Fujifilm Wako Pure Chemical) and incubated in PBS containing 0.1% Tween-20 (T-PBS) supplemented with 5% skimmed milk. After washing with T-PBS, the membranes were incubated with the following primary antibodies: anti-Daed (1:1,000 dilution, this study), anti-Gasz (1:1,000 dilution),^12^ anti-Armi (2F8A9, 1:1,000 dilution),^16^ anti-Piwi (3G11, 1:1,000 dilution),^36^ anti-β-Tub (1:1,000 dilution, E7, Developmental Studies Hybridoma Bank), anti-HSP60(1:500 dilution, LK1, StressMarq Biosciences), anti-Flag M2 (1:5,000 dilution, F3165, Sigma), anti-Histon H3 (1:5,000 dilution, ab1791, abcam) monoclonal and anti-c-Myc (1:1,000 dilution, C3956, Sigma) polycronal antibodies. After washing with T-PBS, the membranes were incubated with peroxidase-conjugated anti-mouse IgG antibodies (1:5,000 dilution, 55558, Cappel) or anti-rabbit IgG antibodies (1:5,000 dilution, 7074, Cell Signaling Technology). The membranes were then reacted with Clarity Western ECL Substrate (Bio-Rad) and images were collected using ChemiDoc XRS Plus System (Bio-Rad). The signal intensity was quantified by Fiji (Ver. 2.0.0-rc-69/1.52p).^71^

### Immunofluorescence

Immunofluorescence was performed as described previously.^16^ OSCs were fixed with 4% formaldehyde in PBS. The cells were permeabilized with PBS containing 0.1% Triton X-100, blocked with PBS containing 3% BSA, and incubated with the following primary antibodies: anti-Armi (2F8A9, 1:200 dilution),^16^ anti-Piwi (4D2, 1:200 dilution,^36^ anti-Yb (1G2H3, 1:200 dilution),^56^ anti-Flag M2 (F3165, 1:1,000 dilution, Sigma) monoclonal and anti-c-Myc (1:300 dilution, C3956, Sigma) polyclonal antibodies. Anti-Flag polyclonal antibodies (F7425, 1:300 dilution, Sigma) were used to obtain the results shown in Figure 2F. After washing with PBS, cells were incubated with the following secondary antibodies (1:1,000 dilution): Alexa 488-conjugated anti-mouse IgG1, Alexa 488-conjugated anti-mouse IgG2b, Alexa 555-conjugated anti-mouse IgG1, Alexa 555-conjugated anti-mouse IgG2b, Alexa 488-conjugated anti-rabbit IgG and Alexa 633-conjugated anti-rabbit IgG (all from Thermo Fisher Scientific). Mitochondria were stained with MitoTracker Orange CMXRos (Thermo Fisher Scientific) before fixation. Cells were mounted with VECTASHIELD Mounting Medium with DAPI (Vector Laboratories). Images were obtained using LSM980 laser scanning confocal microscope (Carl Zeiss).

### Immunoprecipitation

Immunoprecipitation was performed as described previously.^16^ Cells were suspended in binding buffer A [30 mM HEPES-KOH (pH 7.3), 150 mM KOAc, 5 mM Mg(OAc)_2_, 5 mM DTT, 1% Triton X-100, 2 μg/mL leupeptin, 2 μg/mL pepstatin A, and 0.5% aprotinin] supplemented with 0.04 U/μL RNasin Plus RNase Inhibitor (Promega) and the cell suspension was passed 10 times through a 30-gauge needle and centrifuged. The obtained supernatants (cell lysates) were incubated at 4°C with Dynabeads Protein G (Thermo Fisher Scientific) pre-bound with antibodies. After 1 h of incubation, the beads were washed with binding buffer and the materials bound on the beads were analyzed.

### Piwi-piRISC isolation from OSCs

Cells were lysed by sonication in binding buffer B [50 mM HEPES-KOH (pH 7.3), 200 mM KCl, 1 mM EDTA, 1 mM DTT, 1% Triton X-100, 0.1% Na deoxycholate, 2 μg/mL leupeptin, 2 μg/mL pepstatin A, and 0.5% aprotinin] supplemented with 0.1 U/μL RNasin Plus RNase Inhibitor. Cell lysates were then incubated with anti-Piwi monoclonal antibodies (3G11) pre-bound to Dynabeads Protein G at 4°C for 2 h. The beads were washed with IP-wash buffer [20 mM HEPES-KOH (pH 7.3), 300 mM NaCl, 1 mM DTT, 0.05% NP-40, 2 μg/mL leupeptin, 2 μg/mL pepstatin A, and 0.5% aprotinin] and with high-salt IP-wash buffer [20 mM HEPES-KOH (pH 7.3), 500 mM NaCl, 1 mM DTT, 0.05% NP-40, 2 μg/mL leupeptin, 2 μg/mL pepstatin A, and 0.5% aprotinin]. RNAs were eluted from the beads with phenol-chloroform and precipitated with ethanol. Eluted RNAs were separated on denaturing polyacrylamide gels and detected by SYBR Gold Nucleic Acid Gel Stain (Thermo Fisher Scientific). Signal intensity was calculated using Fiji, and displayed using Python3 and seaborn (Ver.0.12.2). For piRNA sequencing, cells were lysed in binding buffer C [20 mM HEPES-KOH (pH 7.3), 150 mM NaCl, 1mM EDTA, 1 mM DTT, 0.5% NP-40, 2 μg/mL leupeptin, 2 μg/mL pepstatin A, and 0.5% aprotinin] supplemented with 0.04 U/μL RNasin Plus RNase Inhibitor. RNAs were separated on denaturing polyacrylamide gels and 20–40-nt RNAs were eluted from the gels and cloned as described previously.^8^ QIAseq FastSelect-rRNA Fly Kit (QIAGEN) was used to remove rRNA from the library.

### Northern blotting

Total RNAs were isolated from OSCs using ISOGEN II (Fujifilm Wako Pure Chemical) and treated with 0.04 U/μL TURBO DNase (Life Technologies). Northern blotting was carried out as described previously.^89^ Total RNAs were separated on denaturing polyacrylamide gels, blotted on Hybond-N+ hybridization membranes (Cytiva), and UV cross-linked. The membranes were incubated at 42°C for 30 min in hybridization buffer [200 mM sodium phosphate buffer (pH 7.2), 7% SDS, and 1 mM EDTA] and probed with ^32^P-labeled oligonucleotides at 42°C overnight. The membranes were washed with NB-wash buffer [30 mM sodium citrate (pH 7.0), 300 mM NaCl, and 0.1% SDS] at 42°C and scanned using Typhoon FLA 9500 (Cytiva). Sequences of oligodeoxynucleotides are shown in Table S3.

### RT-qPCR

Total RNAs from OSCs were reverse-transcribed using ReverTra Ace (Toyobo). Resulting cDNAs were amplified with StepOnePlus (Thermo Fisher Scientific) using PowerUp SYBR Green Master Mix (Thermo Fisher Scientific). Sequences of primers are shown in Table S4.

### Recombinant proteins

GST-Daed ΔTM-His was expressed in *E. coli* (Rosetta DE3) at 16°C overnight. *E. coli* was harvested and lysed by sonication in lysis buffer [30 mM Tris-HCl (pH 8.0), 500 mM NaCl, 1% Triton X-100, 10 mM imidazole, 1× cOmplete ULTRA Tablets (Roche), and 1 mM DTT]. The supernatants after centrifugation were bound to Ni-Sepharose 6 Fast Flow (Cytiva) at 4°C for 1 h. After washing with wash buffer A [30 mM Tris-HCl (pH 8.0), 500 mM NaCl, 1% Triton X-100, 10 mM imidazole, and 1 mM DTT], proteins were eluted from the beads with elution buffer A [30 mM Tris-HCl (pH 8.0), 150 mM NaCl, 1% Triton X-100, 250 mM imidazole, and 1 mM DTT]. Eluted proteins were bound to Glutathione Sepharose 4B (Cytiva) at 4°C for 1 h. After washing with wash buffer B [30 mM Tris-HCl (pH 8.0), 500 mM NaCl, 1% Triton X-100, and 1 mM DTT], proteins were eluted from the beads with elution buffer B [30 mM Tris-HCl (pH 8.0), 150 mM NaCl, 1% Triton X-100, 1 mM DTT, and 15 mM glutathione], dialyzed in dialysis buffer [30 mM Tris-HCl (pH 8.0), 150 mM NaCl, 1 mM DTT, 10% glycerol, and 8 μg/mL GST-HRV3C protease], and passed through Glutathione Sepharose 4B to remove the protease and GST.

### Pull-down assay

OSCs expressing Armi WT-Flag and its ΔN34 mutant were suspended with bait-preparing buffer [30 mM HEPES-KOH (pH 7.3), 150 mM KOAc, 500 mM NaCl, 5 mM Mg(OAc)_2_, 5 mM DTT, 1% Triton X-100, 2 μg/mL leupeptin, 2 μg/mL pepstatin A, 0.5% aprotinin, and 25 μg/mL RNase A (Roche)]. The cell suspension was passed five times through a 25-gauge needle and another five times through a 30-gauge needle, and then centrifuged. The supernatants were incubated at 4°C for 1 h with anti-Flag antibodies pre-bound onto Dynabeads Protein G. Beads were washed with bait-preparing buffer and then binding buffer D [30 mM Tris-HCl (pH 8.0), 150 mM NaCl, 1 mM DTT, and 1% Triton X-100]. Daed ΔTM-His was added to the beads and incubated at 4°C for 1.5 h. After washing with binding buffer, proteins were eluted from the beads, separated on an SDS polyacrylamide gel, and stained with CBB.

### β-elimination assay

β-elimination was performed as described previously with some modifications.^37^ piRNAs were isolated from Piwi immunoprecipitated from OSCs. miRNAs were isolated from Ago1 immunoprecipitated from OSCs using monoclonal anti-Ago1 antibodies^57^. RNAs were incubated in 10 mM NaIO_4_ for 40 min at 0°C in the dark. After ethanol precipitation, RNAs were dissolved in 1 M L(+)-lysine-HCl (pH 8.5) and incubated for 90 min at 45°C. After ethanol precipitation, RNAs were separated on a denaturing polyacrylamide gel and detected by SYBR Gold Nucleic Acid Gel Stain.

### piRNA sequencing and bioinformatic analysis

piRNAs were sequenced using MiSeq (Illumina) with MiSeq Reagent Kit v3 (Illumina). piRNA-seq analysis was performed as described previously^8^ with some modifications. Adapter-trimmed reads of ≥ 18 nt in length were mapped against the *Drosophila* genome (BDGP Release 6 + ISO1 MT/dm6) with FlyBase gene model (Release 6.21) using STAR^64^ (Ver. 2.7.8a). Uniquely mapped reads with up to one mismatch were used for the subsequent analysis. Read length of each biological triplicate was used in the analysis in Figures 4B and S6A. After confirming high correlations between samples (Figures S9A and S9B), reads were merged and used for subsequent analysis. Reads mapped to exons of coding genes (Figures S9A and S9B) were counted with HTSeq^65^ (Ver. 0.9.1) using annotation file obtained from FlyBase and normalized to RPM+1 reads. Spearman’s rho was calculated using R^62^ (Ver. 4.3.1). For transposon annotation (Figures S4B and S4C), the annotation file of repeat sequences was obtained from RepeatMasker (dm6-Apr 2006-RepeatMasker open-4.0.6-Dfam 2.0) and transposons were extracted without distinction between int and LTR. Reads mapped antisense to transposons were counted using HTSeq. For the analysis in Figure S4C, reads were normalized to RPM+1 reads. For the analysis in Figure S4B, transposons which mapped > 5 RPM reads in the sum of WT and ΔSAM were selected, and the mean length of mapped reads for each transposon were calculated. For 1U bias and +1U bias analyses (Figures 4C, 4I, and S6B), the 5′ ends and their downstream sequences of piRNAs (1U bias) or *Drosophila* genome sequences around the 3′ ends (+1U bias) were obtained using bedtools^66^ (Ver. 2.30.0) and SAMtools (Ver. 1.9).^67^ Sequence logos were generated using ggseqlogo.^69^ For the phasing analysis (Figure 4D), 3′ end positions around piRNA 5′ end position were normalized to RPM and showed in heatmaps using RSeQC^70^ (Ver. 2.6.4), deepTools^68^ (Ver. 3.3.2), SAMtools^67^ (Ver. 1.9) and “wigToBigWig” from UCSC. Binary histograms were generated using the matrix of heatmaps. Values were transformed into binary format (counts = 0 → 0, counts > 0 → 1) in accordance with a previous study.^18^ For the comparison of 3′ end position in WT and mutant libraries of Daed or Armi (Figures 4H and S6C), 50,000 piRNA 5′ end positions, common to the WT and mutant libraries and most abundant in the WT library, were extracted. The percentage of 3′ end positions corresponding to each 5′ end position was calculated and shown in heatmaps. 5′ end positions were ordered by the 3′ end position of WT library data and the same order was maintained in the mutant library data. In Figure S4G, the length of dominant piRNAs produced from the 5′ end was used for the 5′-end clarification. First, the 5′ end positions that dominantly produce 26-or 27-nt piRNAs in the WT library was extracted, and after combining, these positions were classified as “Unchanged” and “Elongated” according to the length of dominant piRNAs in ΔSAM library. The subsequent analysis was conducted using the piRNAs produced from each 5′-end groups.

### piRISC production simulation

The *flam* sequence was downloaded from FlyBase (Release FB2023_05) (21,629,891 to 21,792,731 on the X chromosome) and subjected to the digestion into 163 piRNA precursors (starting at U and around 1,000 nt). The precursors were cleaved to 18–40-nt piRNAs immediately before U. The cleavage position was determined based on the distribution of the actual piRNA length, which was obtained by fitting a normal distribution to the piRNA sequencing data. The simulation was performed using Python3^58^ (Ver. 3.8.8) with add-on libraries [Numpy^59^ (Ver. 1.20.1), Scipy^60^ (Ver. 1.6.2), and Pandas^61^ (Ver. 1.2.4)].

### Structure prediction

A structure model of the AmTEC (Figure 1A) and Armi–AmTEC complex (Figure 3A) was built by structure prediction with ColabFold^90^ (Ver. 1.5.5) and AlphaFold^32^ (Ver. 2.3) using protein amino acid sequences from FlyBase (Release FB2023_04). Five complex models were built, and structural predictions and optimization were performed five times for each complex model. The structure model with the highest pLDDT value underwent relax steps. The N-terminal disorder region and TM domain orientations of Daed and Gasz were appropriately changed at the linker region using ISOLDE^74^ (Ver. 1.6). The structure was visualized using UCSF ChimeraX^73^ (Ver. 1.6.1).

### Molecular dynamics simulation

The protein complex was embedded in a bilayer membrane in the simulation box described below using CHARMM-GUI Martini bilayer maker^75–77^ (Ver. 3.8). Orientation of the protein complex in lipid bilayer was determined using PPM 3.0 Web Server^78^ prior to the construction of the simulation box. The structure model of the Armi–AmTEC [Daed (ΔSAM)] complex was generated using PyMOL (2.5.0), UCSF ChimeraX, and ISOLDE based on the Armi–AmTEC complex model. Initial structures were arranged in the 250×250 nm bilayer membrane (67% of POPC and 33% DOPE, containing roughly 2,000 lipid molecules in total) and the thickness of water at the top and bottom of the system was set to 100 nm. Ionic strength and temperature were set to 150 mM NaCl and 303.15 K, respectively. Molecular dynamics simulations were performed using GROMACS^79^ (Version 2021.2) with Martini 3 coarse-grained model.^91^ First, energy minimization was performed using the steepest-descent method. Then, equilibration was conducted using the V-rescale method five times (2 fs×500,000 steps, 5 fs×200,000 steps, 10 fs×100,000 steps, 15 fs×50,000 steps, and 20 fs×50,000 steps). After the equilibration runs, simulations were performed at 20 fs for 50,000,000 steps. The structural trajectories were calculated using gmx trjconv tool for clustering and centering of the protein complex. The root mean square deviation (RMSD) of protein complexes were calculated using gmx rms tool. The distances between the center of mass of Armi helicase domain (Ser697–Cys1,148) and that of AmTEC were calculated using Python3 (Ver. 3.9.18) with add-on libraries [Numpy (Ver. 1.24.3) and MDAnalysis^80,81^ (Ver. 2.7.0)], and visualized using R.^62^

### Ortholog search and protein sequence alignment

Orthologs for Armi/MOV10L1, Gasz/ASZ1, PNLDC1, Papi/TDRKH, Nbr were searched using OrthoDB v11. Orthologs for Daed were searched using OrthoDB v11^39^ and NCBI PSI-Blast. Species taxonomy was according to NCBI and phylogenetic tree (Figure 5A) were generated following previous studies.^92,93^ Multiple sequence alignment (Figure S5B) was conducted using CLUSTAL Omega multiple alignment^82^ and visualized using ESPript (Ver. 3.0).^84^ The phylogenetic tree (Figure S5A) was generated using Neighbor-Joining method^94^ in MEGA11^85^. Multiple sequence alignment (Figure S6) was conducted using CLUSTAL Omega multiple alignment^82^ and visualized using iTOL (Ver. 6.9)^83^. The following protein sequences were acquired from NCBI: *D. melanogaster* Armi (NP_001014556), *H. sapiens* Moloney leukemia virus 10-like protein 1 (MOV10L1) (NP_061868.1), *M. musculus* MOV10L1 (XP_006521619.1), *G. gallus* MOV10L1 (XP_040514825.1), *X. tropicalis* MOV10L1 (NP_001072624.1), *D. rerio* MOV10L1 (NP_001070795.1), *Z. nevadensis* Armi (XP_021937793.1), *T. palmi* Armi (XP_034251738.1), *A. echinatior* Armi (XP_011062616.1), *B. terrestris* Armi (XP_012166280.1), *D. ponderosae* Armi (XP_048525683.1), *T. castaneum* Armi (XP_008199251.1), *B. anynana* Armi (XP_023946786.2), *B. mori* Armi (XP_037867032.1), *A. gambiae* Armi (XP_061506580.1), *A. aegypti* Armi (XP_021698921.1), *B. dorsalis* Armi (XP_011214121.1), *C. capitata* Armi (XP_004537921.1), *M. domestica* Armi (XP_058975516.1) and *S. calcitrans* Armi (XP_013110641.2).

### Quantification and statistical analysis

All experiments were performed over two independent experiments (biological replicates). Sample sizes were predetermined following previously performed experiments with similar setup in the field. Statistical methods, *p* value and number of replicates are indicated in each figure and its legend. Statistical analyses were performed in the R software environment (version 4.3.1). Shapiro-Wilk test was used for testing normality when statistical methods based on the normal distribution was used. Experiments were not blinded as consistent with the previous studies in the field.

## Supplemental information

Document S1. Figures S1–S9 and Table S1–S4

## Supplemental information

**Figure S1. Related to Figure 1.**
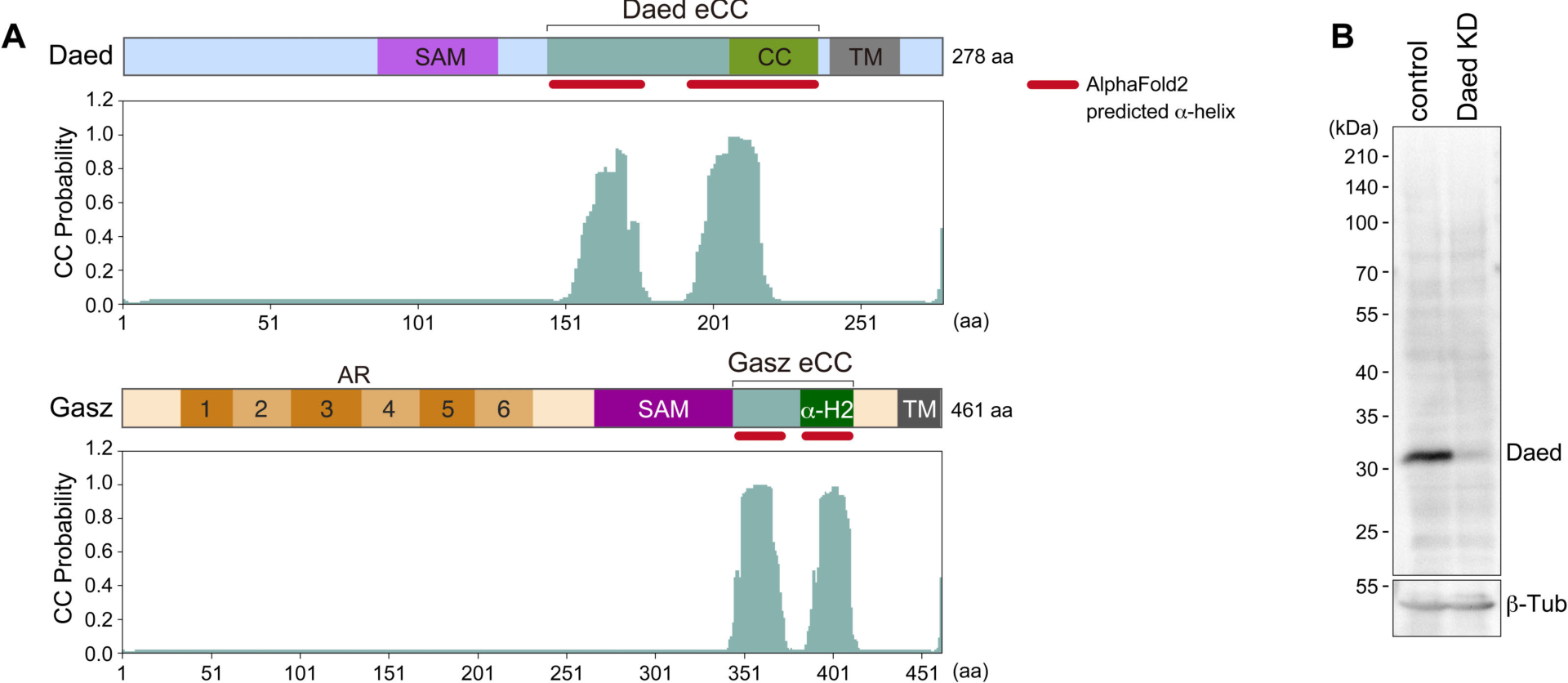
Coiled-coil prediction and specificity of anti-Daed antibodies. (**A**) Sequence-based CC prediction of Daed (upper) and Gasz (lower) by CoCoNat. Red line indicates the CC region predicted by AlphaFold2. (**B**) Western blotting shows that anti-Daed monoclonal antibodies produced in this study are specific to Daed. Daed KD: Daed knockdown by RNAi. control: EGFP KD. β-Tub: loading control.

**Figure S2. Related to Figure 2.**
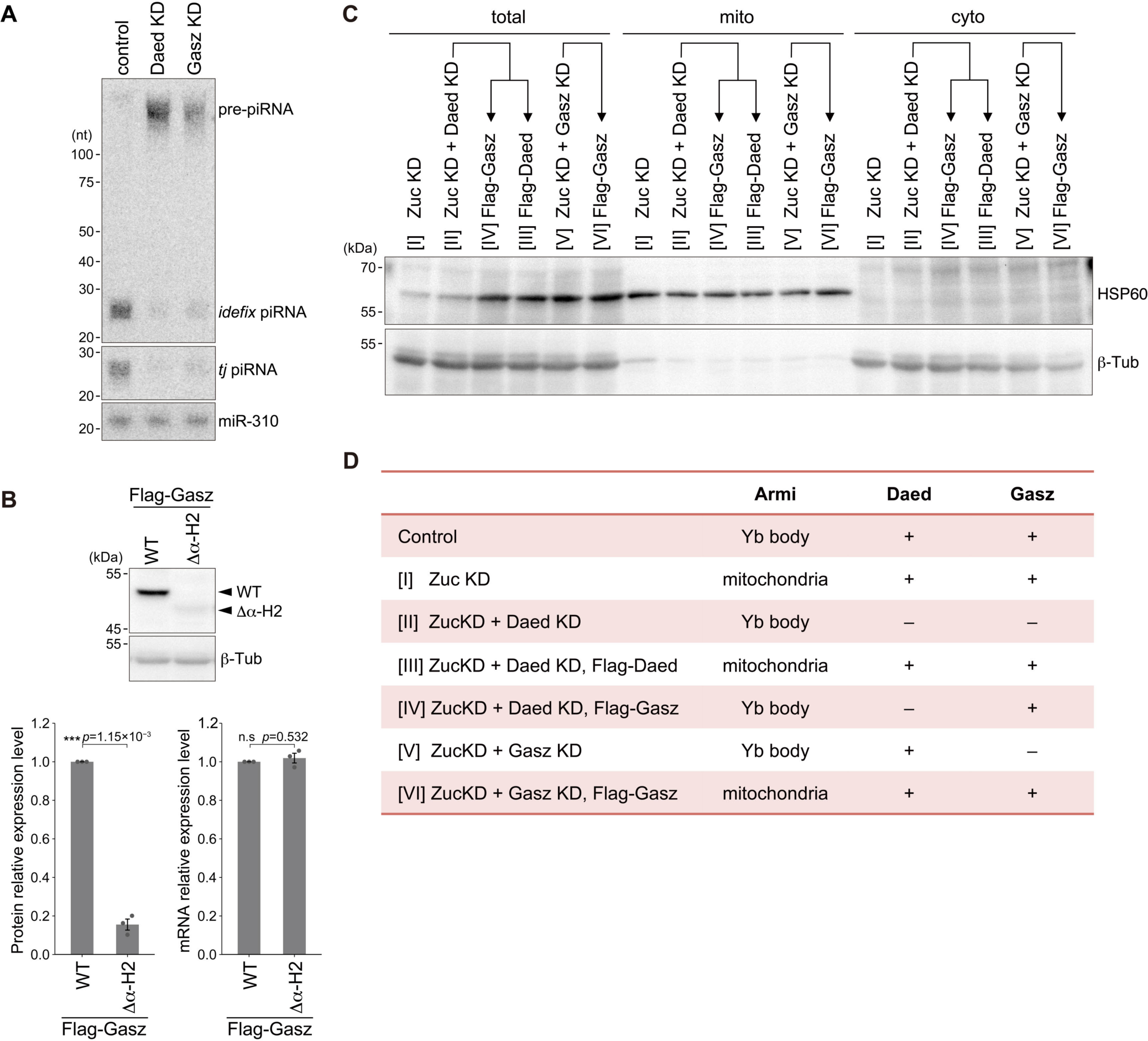
Functions and behaviors of Daed and Gasz in OSCs. (**A**) Northern blotting using probes that detect *idefix* piRNA and *tj* piRNA. miR-310: loading control. KD: knockdown by RNAi. control: EGFP KD. pre-piRNA: piRNA precursors. (**B**) Western blotting shows the levels of Flag-Gasz WT and Flag-Gasz Δα-H2 where the same amounts of plasmids were used for transfection. β-Tub: loading control. The statistics of western blotting signals (n= 3; left) and the mRNA expression levels (n= 3; right). Signal intensities (left) or values (right) relative to F-Gasz WT were normalized with those of β-Tub or *rp49*, respectively, and presented as mean values ± SE. ***: *p* <0.005; n.s.: *p* >0.05 [*t*-test (unpaired, two-sided)]. (**C**) Western blotting shows HSP60 (a mitochondrial fraction marker) and β-Tub (a cytoplasmic fraction marker) in total lysates, and mitochondrial and cytoplasmic fractions in Figure 2G. (**D**) Summary of Armi subcellular localization in OSCs under each condition.

**Figure S3. Related to Figure 3.**
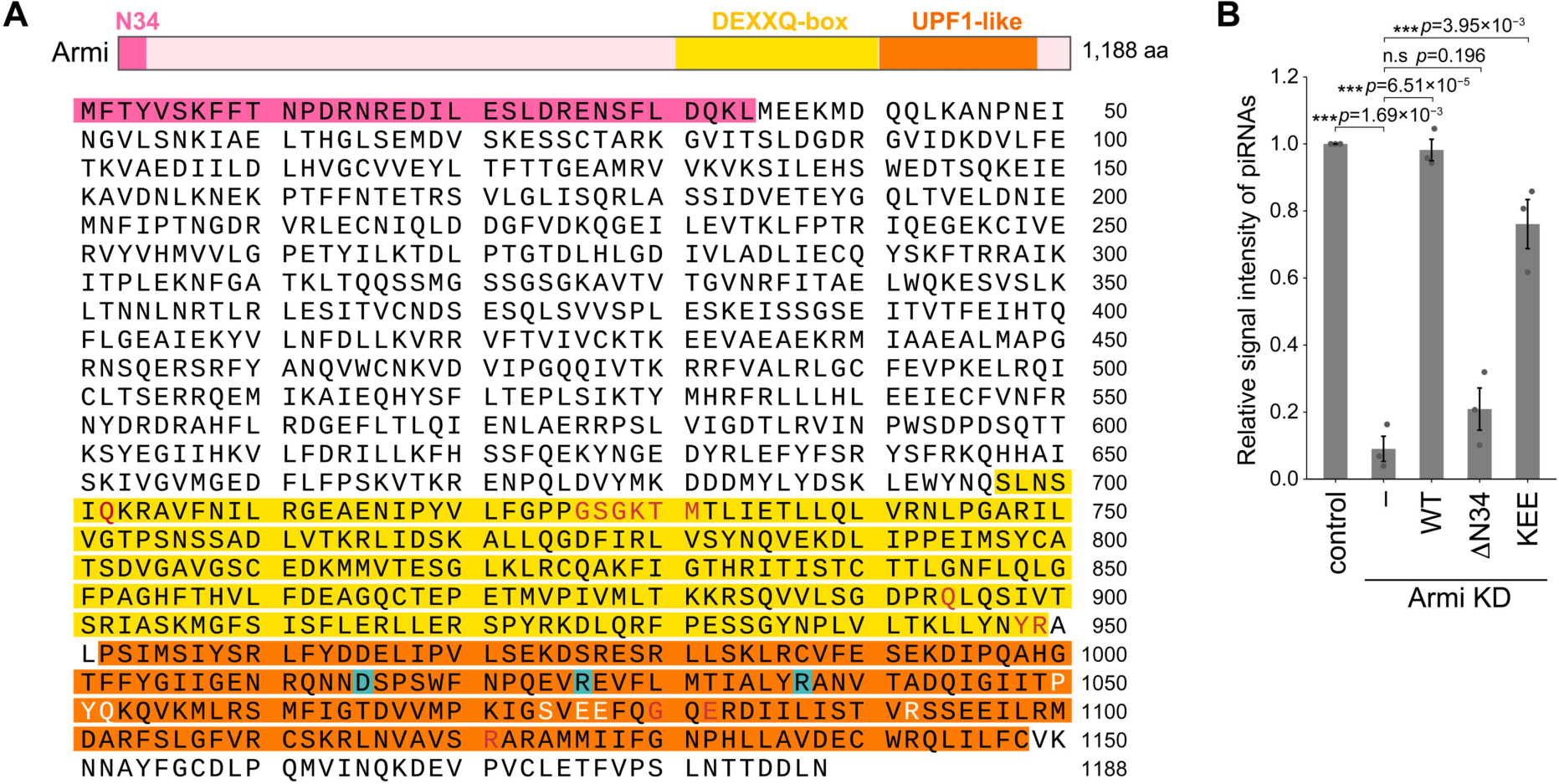
Effects of Armi mutants on piRNA biogenesis. (**A**) Peptide sequence and domain structure of Armi. The ATP binding residues and nucleotide binding residues are indicated in red and white, respectively. Asp1015, Arg1026, and Arg1037 are highlighted in blue. (**B**) Quantification of piRNA levels (n=3) (Figure 3H). Values relative to the control (EGFP KD) were normalized to Piwi level and presented as mean values ± SE. ***: *p* <0.005; n.s.: *p* >0.05 [*t*-test (unpaired, two-sided)].

**Figure S4. Related to Figure 4.**
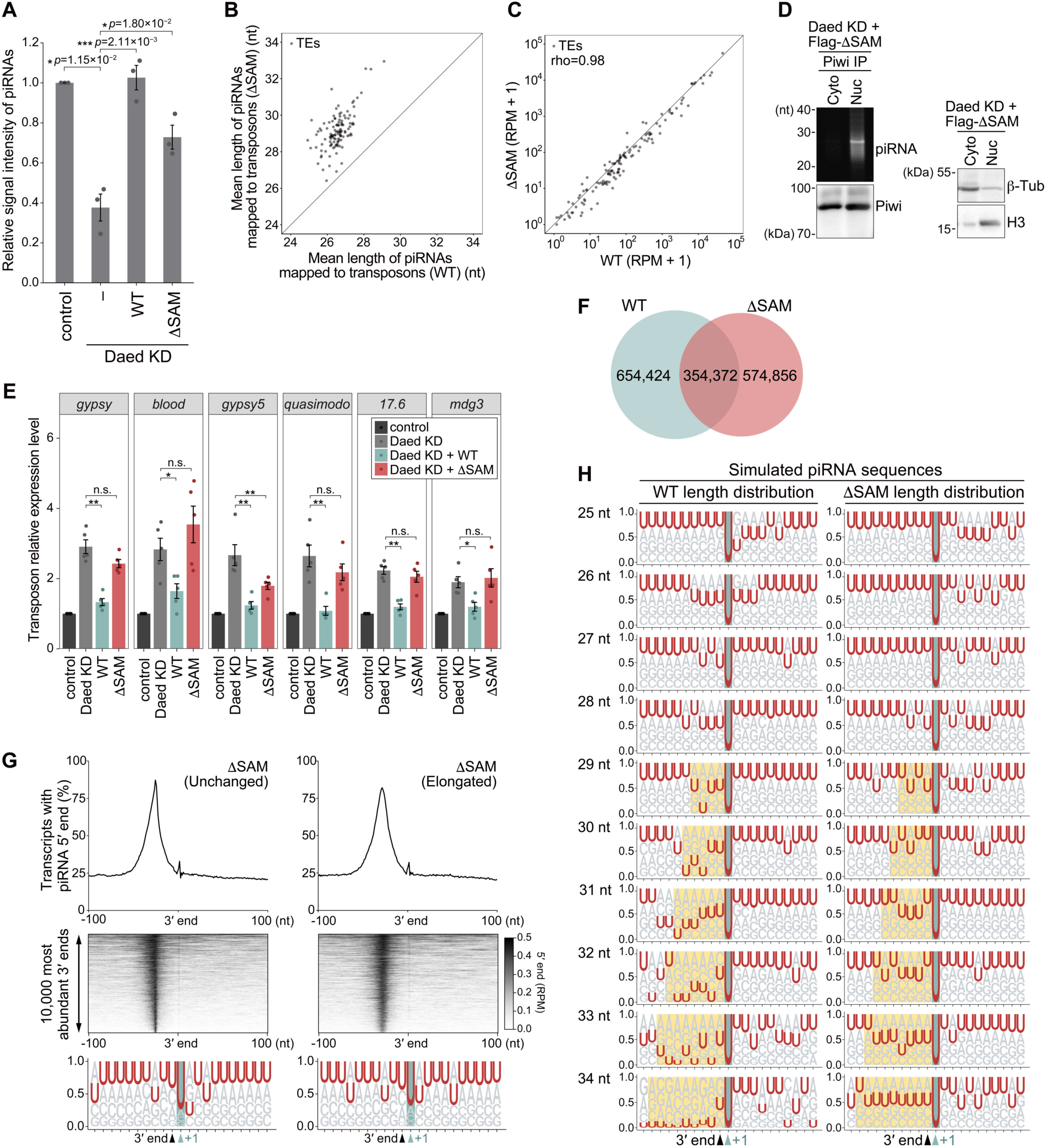
Effects of Daed ΔSAM on piRNA biogenesis and Piwi-piRISC function. (**A**) Quantification of piRNA levels (n=3) (Figure 4A). Values relative to the control (EGFP KD) were normalized to Piwi level and presented as mean values ± SE. ***: *p* <0.005; *: *p* <0.05; [*t*-test (unpaired, two-sided)]. (**B**, **C**) Scatter plots showing the length distribution of piRNAs upon subdivision by target transposons (**B**) and the abundance of reads mapped antisense to transposons (**C**; Spearman’s rho=0.98) in Daed WT and ΔSAM libraries. (**D**) Left: Piwi-bound piRNAs obtained from cytoplasmic fraction (Cyto) and nuclear extract fraction (Nuc). Immunoprecipitated Piwi was detected by western blotting. Right: Western blotting showing β-Tub (a cytoplasmic marker) and Histon H3 (a nucleus marker) in each fraction. (**E**) The expression levels of transposons (*gypsy*, *blood*, *gypsy5*, *quasimodo*, *17.6*, and *mdg3*) in control OSCs (dark gray), Daed-lacking OSCs (Daed KD; light gray), Flag-Daed WT rescued (WT; green) or Flag-Daed ΔSAM rescued cells (ΔSAM; red) (n=5). Values relative to the control (EGFP KD) were normalized to *rp49* and presented as mean values ± SE **: *p* <0.01; *: *p* <0.05; n.s.: *p* >0.05 [Wilcoxon rank sum exact test (two-sided)]. (**F**) Venn diagram showing the variety of positions of piRNA 5′ ends in the WT and ΔSAM libraries. (**G**) Heatmaps showing 5′ end counts of piRNAs mapped around the 10,000 most abundant piRNA 3′ ends (middle) and its binary histogram (top). In sequence logos (bottom), piRNAs were aligned according to the 3′ end of the piRNAs (black arrowhead) and its immediately downstream nucleotide is highlighted in green. Left: Daed ΔSAM piRNAs in “Unchanged” group. Right: Daed ΔSAM piRNAs in “Elongated” group. (**H**) piRNA production simulation. The U depletion in 29–34-nt piRNAs in WT and equivalent regions in ΔSAM are highlighted in yellow.

**Figure S5. Related to Figure 4.**
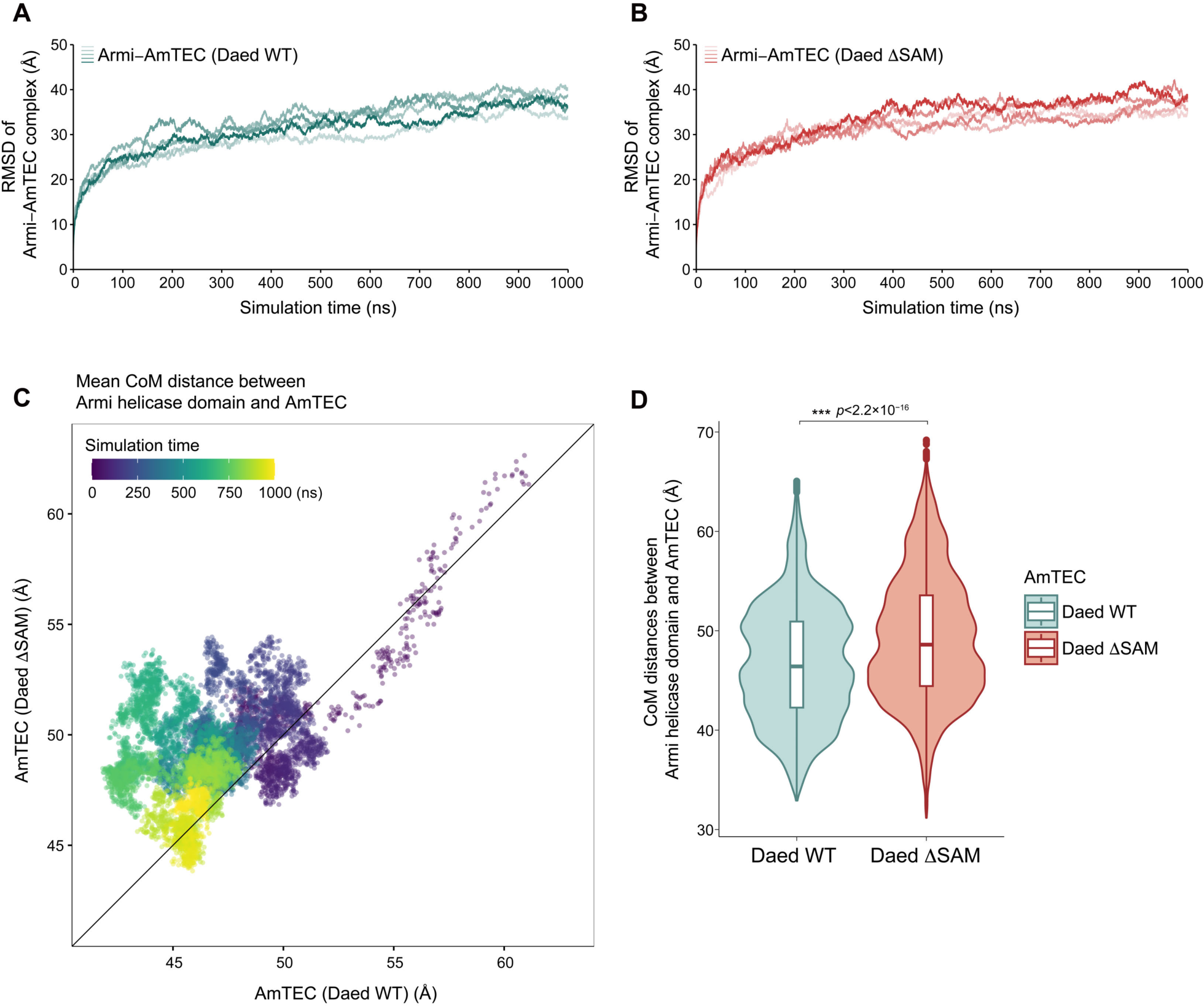
MD simulation of Armi–AmTEC complex. (**A**, **B**) Root mean square deviation (RMSD) of the Armi–AmTEC complex (**A**) and its Daed ΔSAM version (**B**) in molecular dynamics simulations (n=5) indicates that the simulation reaches equilibrium state over the simulation time. (**C**) The average distance between the center of mass (CoM) of Armi helicase domain and that of AmTEC (n=5) for each time point of the trajectory is shown. (**D**) Violin plot of CoM distances between Armi helicase domain and AmTEC (blue) and its Daed ΔSAM version (red) for all time points in all trajectories. ***: *p* <0.005 [Welch’s *t*-test (two-sided)].

**Figure S6. Related to Figure 4.**
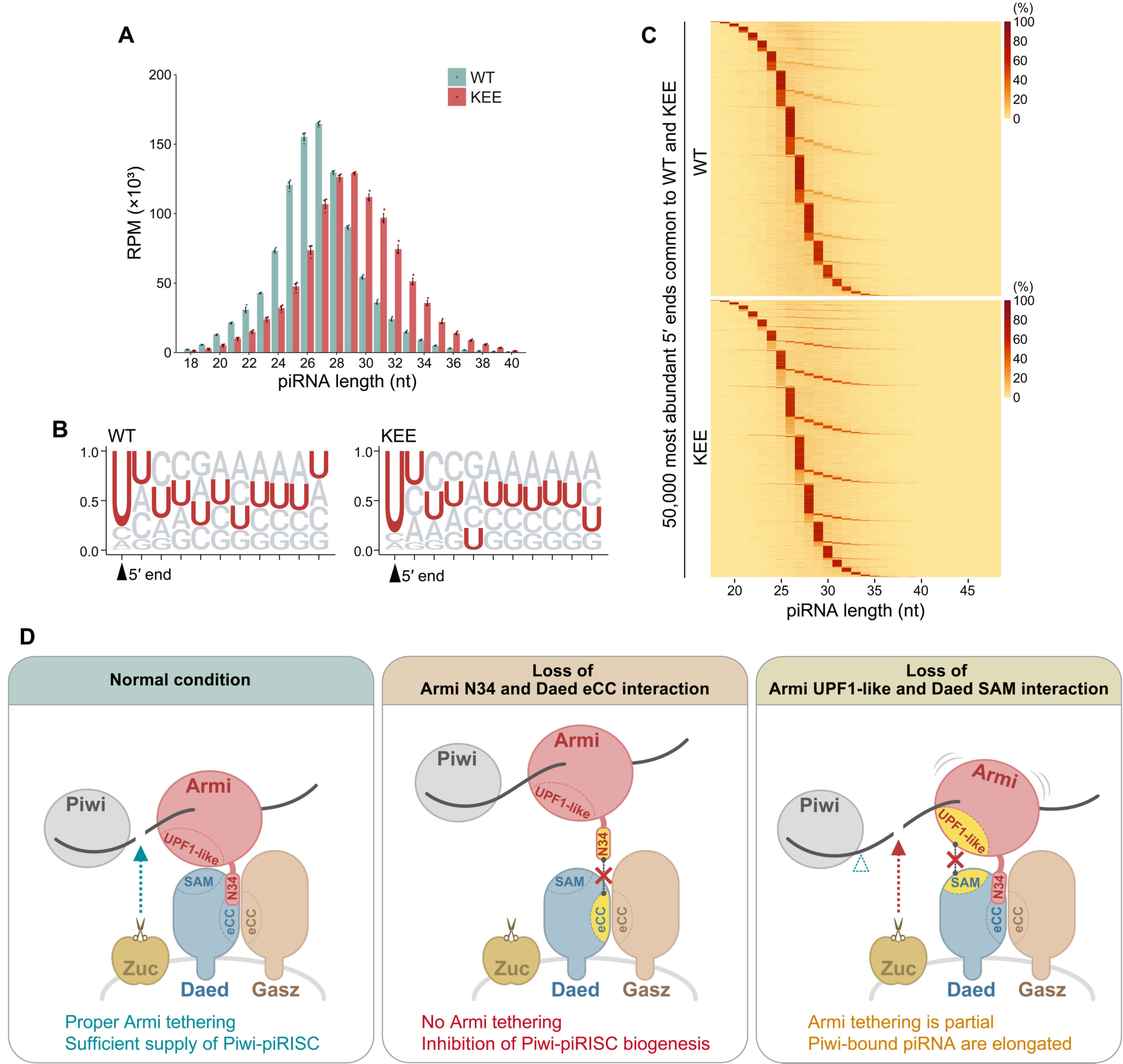
Effects of Armi KEE on piRNA biogenesis. (**A**) Length distribution of piRNAs (18–40 nt) in Armi WT (green) and KEE (red) libraries (n= 3), presented as mean values ± SE. (**B**) Sequence logos representing 11 nt from the 5′ end of piRNAs in the Armi WT and KEE libraries. (**C**) Heatmaps showing the frequency of the 3′ end positions (red) corresponding to each 5′ end of piRNAs in Armi WT (upper) and KEE (lower) libraries. The 50,000 piRNA 5′ end positions common to both libraries and most abundant in the WT library were used for analysis. (**D**) The model summarizing the effects of the perturbation of two Armi–AmTEC interactions (Armi N34–Daed eCC and Armi UPF1-like–Daed SAM). Blue arrowhead (left panel): Zuc cleavage. Red arrowhead (right panel): obscured Zuc cleavage. White arrowhead (right panel): Zuc cleavage corresponding to WT. Black dashed line with a red cross (middle and right panel): perturbated interaction.

**Figure S7. Related to Figure 5.**
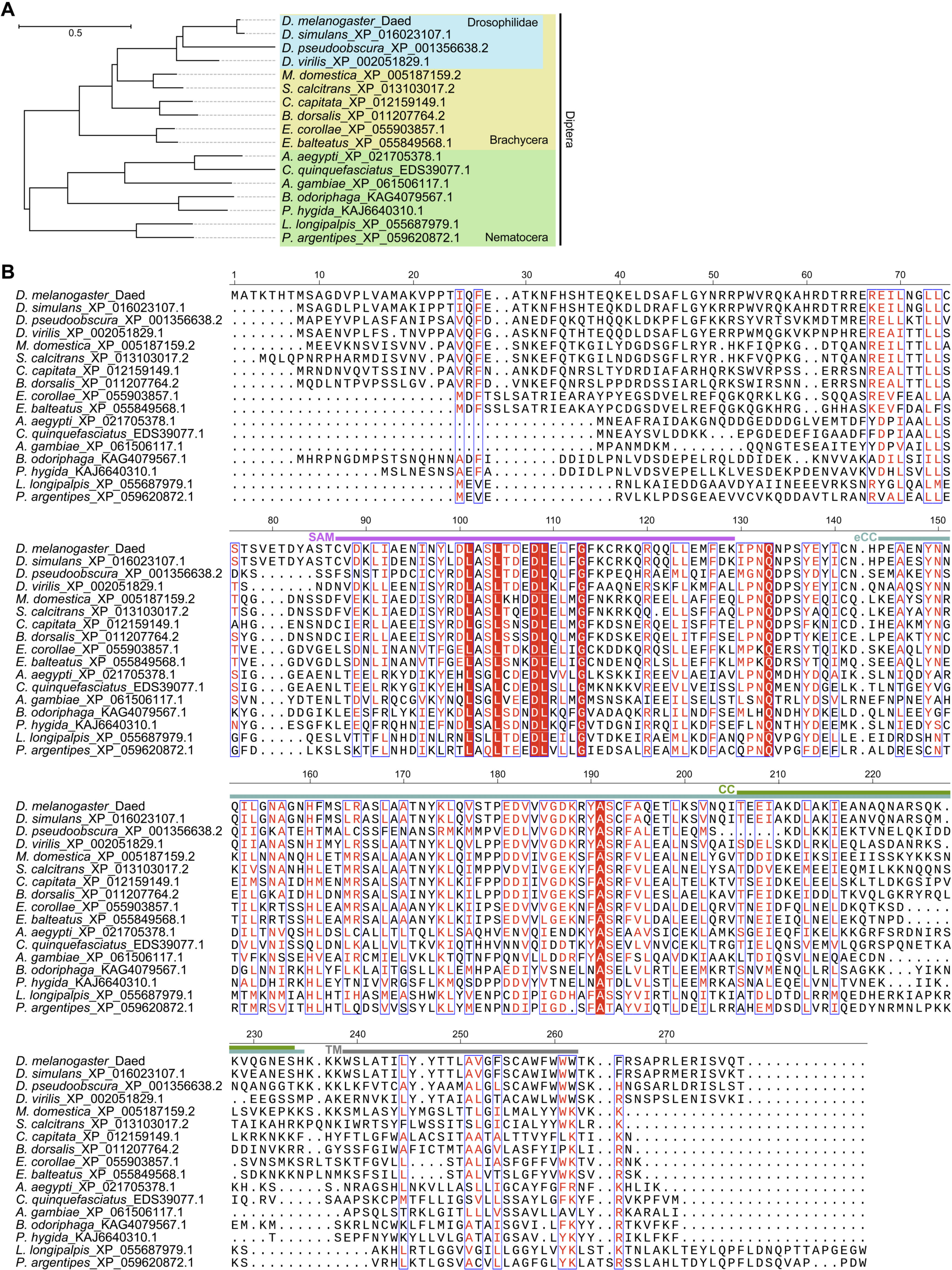
Sequence alignment of Daed in species. (**A**, **B**) Phylogenetic tree (**A**) and multiple sequence alignment (**B**) of Daed orthologs constructed and visualized using Clustal Omega, MEGA 11 and ESPript.

**Figure S8. Related to Figure 5.**
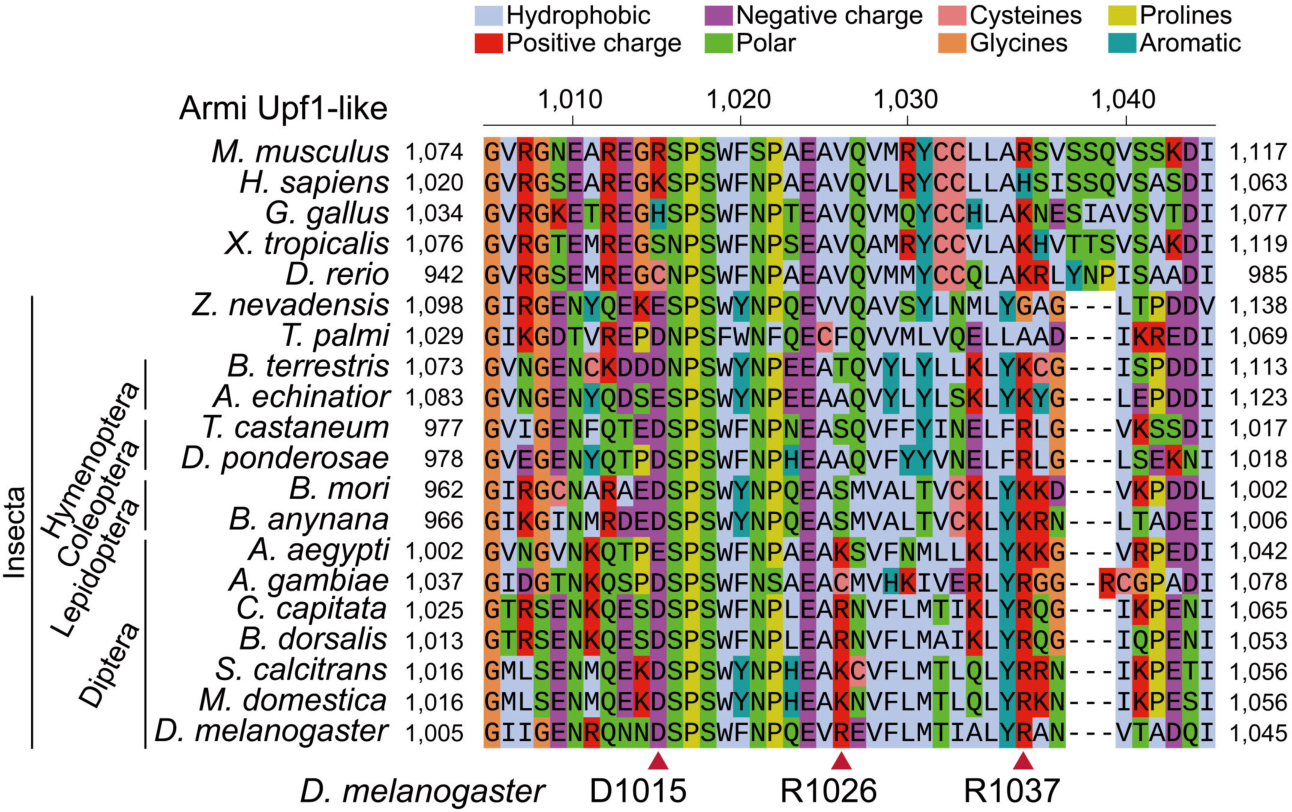
Sequence alignment of Armi UPF1-like helicase domain in species. Multiple sequence alignment of *D. melanogaster* Armi and its orthologs. Red arrowheads: Asp1015, Arg1026, and Arg1037 (D-R-R) of *D. melanogaster* Armi.

**Figure S9.**
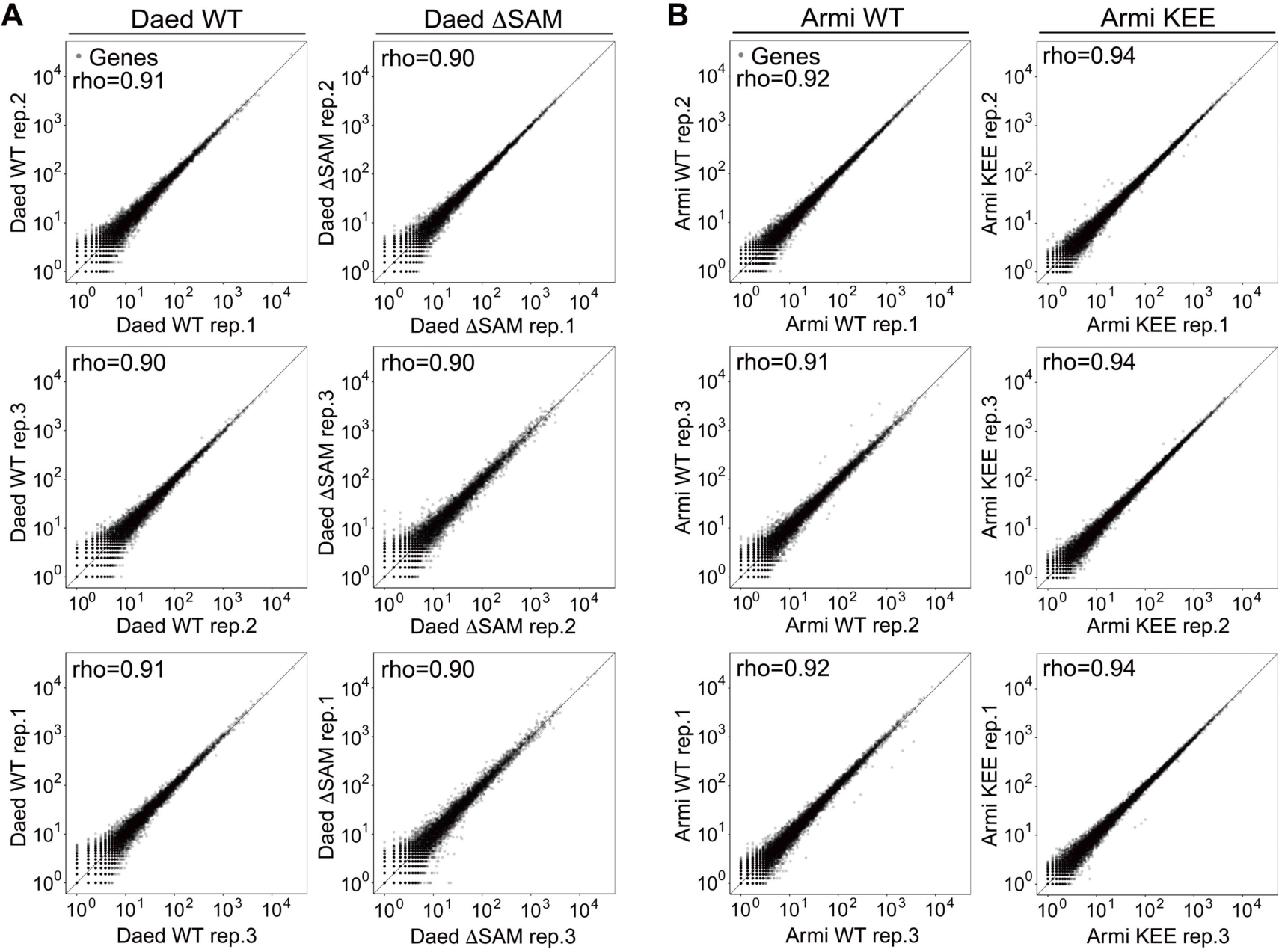
Related to STAR Method “piRNA sequencing and bioinformatic analysis”; Reproducibility of piRNA-seq. (**A,B**) Scatter plots showing reproducibility of piRNA-seq of Daed WT, ΔSAM mutant (**A**), Armi WT and KEE mutant libraries (**B**). Gray dots indicate the number of normalized piRNA reads mapped to coding genes.

**Table S1.**
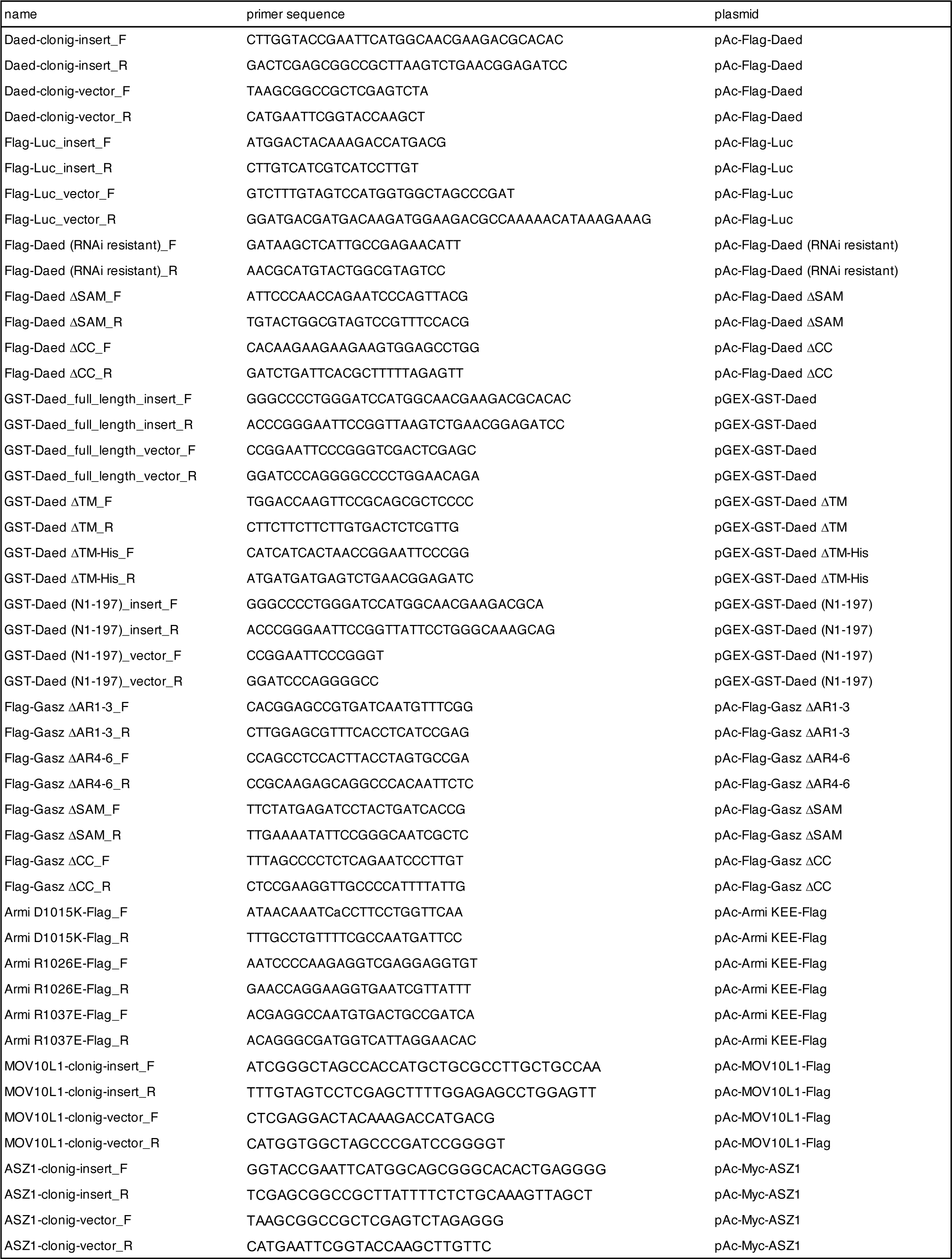
Related to STAR Method “Plasmid construction”; Sequences of primers used for plasmid construction.

**Table S2.**
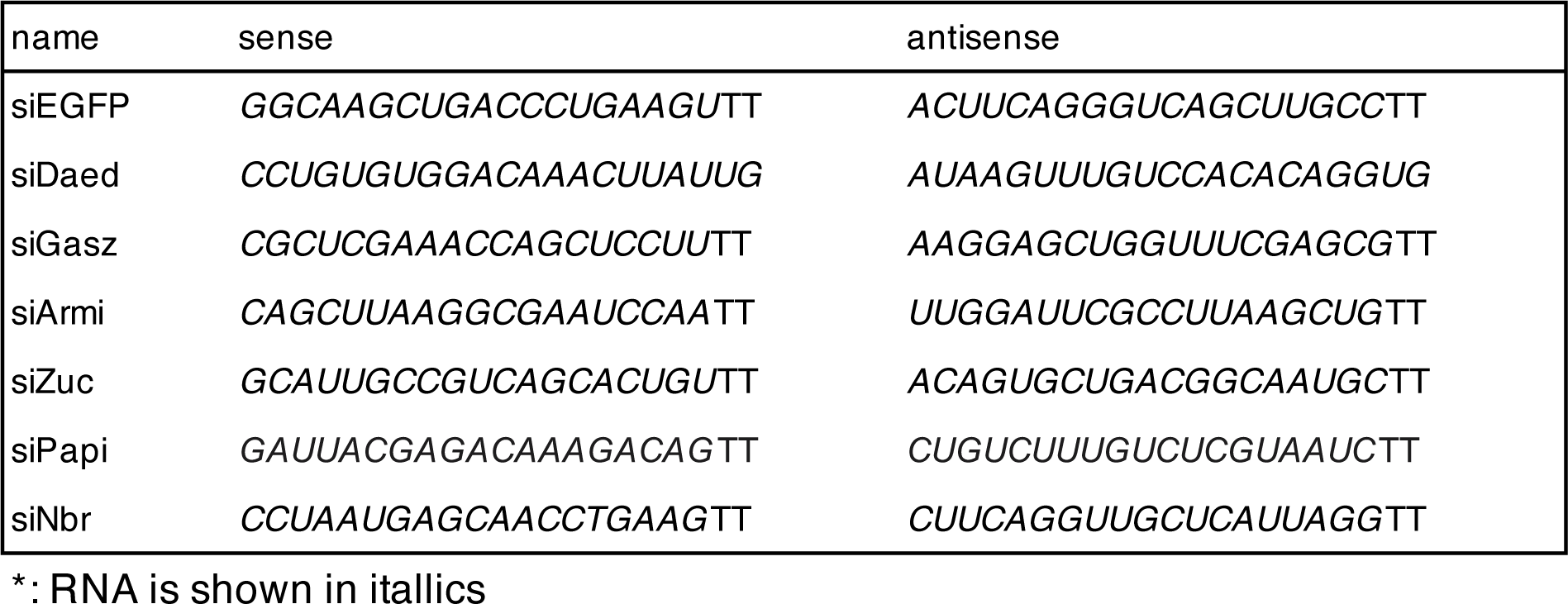
Related to STAR Method “RNAi and transfection”; Sequences of siRNAs.

**Table S3.**
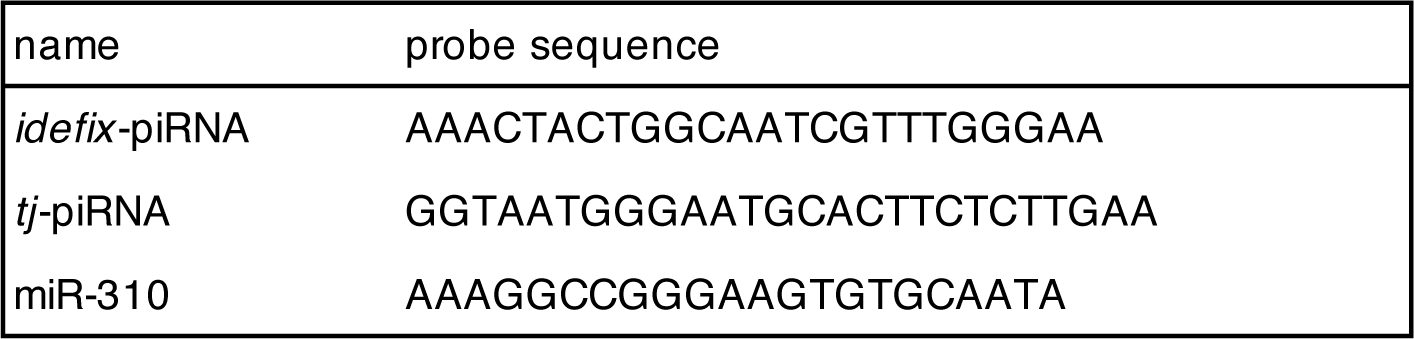
Related to STAR Method “Northern blotting”; Sequences of oligos used for northern blotting.

**Table S4.**
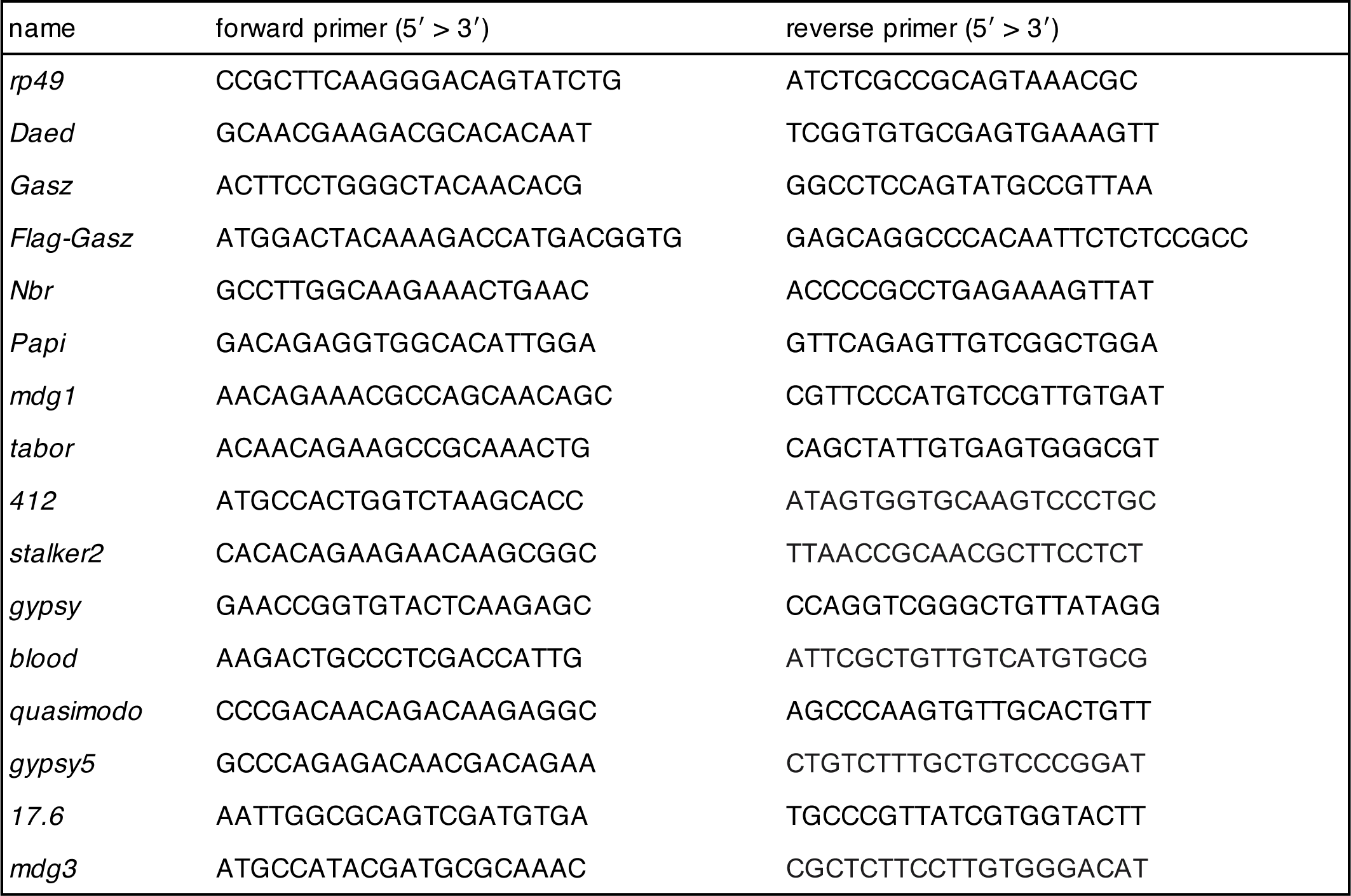
Related to STAR Method “RT-qPCR”; Sequences of primers used for RT-qPCR.

